# ARL3 Mediates BBSome Ciliary Turnover by Promoting Its Outward Diffusion through the Transition Zone

**DOI:** 10.1101/2021.11.19.469163

**Authors:** Yan-Xia Liu, Wei-Yue Sun, Bin Xue, Rui-Kai Zhang, Wen-Juan Li, Xixian Xie, Zhen-Chuan Fan

**Author notes:** All the correspondence should be addressed to: Zhen-Chuan Fan, Ph.D. Rm 334, Yifu Building, No. 9 13 St. Tianjin Economic and Technological Development Area Tianjin University of Science and Technology, Tianjin 300457, China.

## Abstract

Ciliary receptors and their certain downstream signaling components undergo intraflagellar transport (IFT) as BBSome cargoes to maintain their ciliary dynamics for sensing and transducing extracellular stimuli inside the cell. Cargo laden BBSomes shed from retrograde IFT at the proximal ciliary region above the transition zone (TZ) followed by diffusing through the TZ for ciliary retrieval, while how the BBSome barrier passage is controlled remains elusive. Here, we show that the BBSome is a major effector of the Arf-like 3 (ARL3) GTPase in *Chlamydomonas*. Under physiological condition, ARL3^GDP^ binds the membrane for diffusing into and residing in cilia. Following a nucleotide conversion, ARL3^GTP^ dissociates with the ciliary membrane and binds and recruits the IFT-detached and cargo (phospholipase D, PLD)-laden BBSome at the proximal ciliary region to diffuse through the TZ and out of cilia. ARL3 deficiency impairs ciliary signaling, e.g. phototaxis of *Chlamydomonas* cells, by disrupting BBSome ciliary retrieval, providing a mechanistic understanding behind BBSome ciliary turnover required for ciliary signaling.

## Introduction

Cilia and flagella are interchangeable terms referring to the axonemal microtube-based subcellular organelles projecting from the cell surface of most eukaryotic cells. They act as antennas for sensing and transducing the extracellular stimuli into the cell, thus essential for maintaining many physiological and developmental signaling pathways (Goetz and Anderson, 2010; Nachury and Mick, 2019; Singla and Reiter, 2006). Therefore, ciliary malfunction causes a group of related genetic disorders including Joubert syndrome, Meckel-Gruber syndrome, nephronophthisis, and Bardet-Biedl syndrome (BBS), collectively named ciliopathies (Hildebrandt et al., 2011). Underlying ciliopathies is the fact that many G protein-coupled receptors (GPCRs), ion channels, and enzymes like receptor tyrosine kinases, platelet-derived growth factor receptor alpha, and insulin-like growth factor-1 position to and traffic inside the ciliary membrane by motor protein-driven intraflagellar transport (IFT) trains along the axoneme (Liu et al., 2020; Nachury and Mick, 2019; Schneider et al., 2005; Yeh et al., 2013). During this process, the BBSome composed of multiple BBS proteins links these ciliary transmembrane signaling proteins to IFT trains composed of repeating units of IFT-A (6 subunits) and -B (16 subunits subdivided into IFT-B1 and -B2 entities) complexes by acting as an IFT cargo adaptor (Fan et al., 2010; Jin et al., 2010; Lechtreck et al., 2009; Loktev et al., 2008; Nachury et al., 2007; Nakayama and Katoh, 2020; Taschner and Lorentzen, 2016; Wang et al., 2009). Compared with these ciliary receptors, channels, and enzymes, their downstream lipidated signaling cascade factors [e.g. heterotrimeric G protein transducin (Gα,β,γ), inositol polyphosphate-5-phosphatase (INPP5E), nephronophthisis 3 (NPHP3)] do not count on the IFT/BBS system for shuttling into cilia to bind the ciliary membrane. They instead use the ADP-ribosylation factor (Arf)-like 3 (ARL3) pathway to achieve this goal.

As a member of the Arf subfamily of the Ras superfamily of small GTPases, ARL3 is conserved among the ciliated species but absent from the non-ciliated organisms, localizes throughout the cell, and is enriched in cilia (Avidor-Reiss et al., 2004; Efimenko et al., 2005; Pazour et al., 2005). Vertebrate and human ARL3 has three types of effectors including phosphodiesterase 6 delta subunit (PDE6D), uncoordinated-119A/B (UNC119A/B), and binder of Arl2 (BART)/binder of Arl2-like 1 (BARTL1), acting in general as carrier/solubilizing proteins to bind and shuttle the cytoplasmic lipidated cargoes of different groups into cilia (ElMaghloob et al., 2021; Linari et al., 1999; Lokaj et al., 2015; Wright et al., 2011). It was known that PDE6D binds and transports the C-terminal prenylated (farnesylated or geranylgeranylated) cargoes [e.g. the catalytic α and β subunits of PDE6, INPP5E, transducin γ subunit (Tγ), and rhodopsin kinase (GRK1)] into cilia (Li and Baehr, 1998; Thomas et al., 2014; Zhang et al., 2007; Zhang et al., 2004). UNC119A/B instead binds and transports the N-terminal myristoylated cargoes [e.g. NPHP3, cystin, and transducin α subunits (GNAT-1 and GNAT-2)] into cilia (Wright et al., 2011; Zhang et al., 2011a). Compared with PDE6D and UNC119A/B, BART/BARTL1 has no cargoes determined thus far, while BART was recently identified to act as a ARL3-specific co-guanine nucleotide exchange factor (GEF) to contribute to convert GDP-bound ARL3 (ARL3^GDP^) to GTP-bound ARL3 (ARL3^GTP^) (ElMaghloob et al., 2021). During cargo ciliary targeting, the carrier protein first binds the lipidated cargo in the cytoplasm and the carrier-cargo complex then shuttles towards cilia by an unknown mechanism. Upon arriving at the transition zone (TZ) region, the activated ARL3 (ARL3^GTP^) binds its carrier protein effector at a site allosterically different from the one for lipidated cargo binding, and this induces conformational changes of the carrier protein, leading to the release of the cargo for binding to the ciliary membrane (Fansa et al., 2016; Ismail et al., 2012; Watzlich et al., 2013). After this, ARL3^GTP^ (the activated form) is bound to and stimulated by retinitis pigmentosa 2 (RP2), the ARL3-specific GTPase activating protein (GAP), to hydrolyze for releasing the carrier protein from ARL3 (Veltel et al., 2008). ARL3^GDP^ (the inactivated form) is then reactivated by the ARL3-specific GEF, ARL13b, and ARL3^GTP^ recycles back to bind the carrier-cargo complex for cargo releasing in cilia (Gotthardt et al., 2015; Zhang et al., 2016).

Besides its role in releasing a variety of lipidated signaling factors for ciliary membrane binding, ARL3 was observed to be essential for mouse rhodopsin and worm and human polycystin-1 and -2 (PKD1 and PKD2) to target to cilia (Schrick et al., 2006; Su et al., 2014; Zhang et al., 2013). Given that these ciliary transmembrane signaling proteins cycle through cilia by IFT via binding the BBSome, ARL3 could mediate their ciliary dynamics through the BBSome (Abd-El-Barr et al., 2007; Liu et al., 2020; Nachury, 2018; Nishimura et al., 2004; Su et al., 2014). Our previous study has shown that Rab-like 5 (RABL5) GTPase IFT22 coordinates with ARL6/BBS3 for recruiting the BBSome, as a BBS3 effector, to the basal body in a GTP-dependent manner in *Chlamydomonas reinhardtii* (Sun et al., 2021; Xue et al., 2020). BBS3 itself also diffuses into cilia and binds the ciliary membrane via its N-terminal amphipathic helix in a GTP-dependent manner (Gillingham and Munro, 2007; Liu et al., 2021). At the ciliary tip, BBS3 binds and recruits the BBSome to the ciliary membrane in a GTP-dependent manner, making it spatially available to couple with the membrane-anchored signaling proteins, e.g. phospholipase D (PLD), for ciliary exit by IFT (Liu et al., 2021; Sun et al., 2021). Prior to this event, IFT/BBS remodels to allow for the BBSome to undergo a disassembly/reassembly cycle (Sun et al., 2021). During this cycle, the heterodimer IFT25/27 composed of IFT25 and RABL4/IFT27 is indispensable for the disassembled BBSome subunits to reassemble at the ciliary tip (Dong et al., 2017; Sun et al., 2021). Interestingly, mammalian GPCRs undergo retrograde IFT from the ciliary tip to a proximal ciliary region right above the TZ (Nachury, 2018; Ye et al., 2018). At this region, the GPCR-loaded BBSome sheds from IFT and diffuses through the TZ for ciliary retrieval in a RABL2-dependent manner (Dateyama et al., 2019; Duan et al., 2021; Nachury, 2018; Ye et al., 2018). However, the molecular mechanism underlying how active transport of the BBSome across the TZ diffusion barrier is controlled remains elusive thus far (Nozaki et al., 2019; Ye et al., 2018).

In this study, we identified the BBSome acts as a major ARL3 effector at the ciliary base right above the TZ but not in the cell body of *C. reinhardtii*. ARL3 mimics other Arf-like GTPases for diffusing into cilia and reversibly binding the ciliary membrane via its N-terminal amphipathic helix and the G2 residue but in a GDP-dependent manner (Liu et al., 2021). Once in cilia, ARL3 undergoes GTPase cycling and the activated ARL3^GTP^ binds the retrograde IFT-detached and PLD-laden BBSome at the proximal ciliary region right above the TZ and recruits it to diffuse through the TZ for ciliary retrieval. Since disruption of BBSome ciliary dynamics generates cell defective in phototaxis, our finding thus fills a gap in our understanding of how ARL3 mediates phototaxis through controlling BBSome ciliary turnover in *C. reinhardtii* (Sun et al., 2021). Our data also shed lights on the molecular mechanism of how ARL3 deficiency could cause BBS disorder in humans.

## Results

### ARL3 diffuses into cilia

*Chlamydomonas* ARL3 shares significant homology with its orthologues in ciliated species and is more closely related to homologues of worms, *leishmania*, and *Trypanosoma* than mammals and humans phylogenetically (Fig. S1 A and B). ARL3 was shown to be a negative regulator of ciliation in *leishmania* and mouse (Cuvillier et al., 2000; Efimenko et al., 2005; Hanke-Gogokhia et al., 2016). In worms, depletion of ARL3 causes IFT-B and KIF17 motor to dissociate through histone deacetylatase 6 (HDAC6)-dependent pathway and then disrupts IFT (Li et al., 2010; Zhang et al., 2013). To clarify whether ARL3 affects IFT and ciliation in *C. reinhardtii*, we examined the ARL3 CLiP mutant (LMJ.RY0420.182282) that we named *arl3*. The *arl3* cell contains a 2,217-bp paromomycin gene insertion in the fourth exon of the ARL3 gene (Fig. S2 A-C). With the newly developed ARL3 antibody available, this insertion was verified to prevent ARL3 from being synthesized, as shown by immunoblotting, demonstrating that *arl3* is a ARL3-null mutant (Fig. S3 A). *arl3* cells grown cilia of normal length, excluding ARL3 from mediating ciliation (Fig. S3 B). Supportive of this conclusion, *arl3* cells retained IFT-A subunits IFT43 and IFT139, IFT-B1 subunits IFT22 and IFT70, and IFT-B2 subunits IFT38 and IFT57 at wild-type (WT) levels both in whole cell and ciliary samples in the steady state (Fig. 1 A). To examine if ARL3 affects IFT ciliary dynamics, we generated transgenic strains *arl3*::IFT43::HA::YFP, *arl3*::IFT22::HA::YFP, and *arl3*::IFT38::YFP, which expresses IFT43, IFT22, or IFT38 fused at their C-terminus to hemagglutinin (HA) and/or yellow fluorescent protein (YFP) (IFT43::HA::YFP, IFT22::HA::YFP, and IFT38::YFP) in *arl3* cells. When expressed at the same level as when three HA::YFP/YFP-tagged proteins of different IFT subcomplexes were expressed alone in CC-125 control cells (resulting strains CC-125::IFT43::HA::YFP, CC-125::IFT22::HA::YFP, and CC-125::IFT38::YFP) (Fig. S3 C) (Xue et al., 2020), they entered cilia (Fig. 1 B) and underwent typical bidirectional IFT of *C. reinhardtii* as reflected by total internal reflection fluorescence (TIRF) assays, excluding ARL3 from mediating IFT ciliary dynamics (Fig. 1 C and D) (Xue et al., 2020). After knowing this, we asked whether and how *Chlamydomonas* ARL3 enters cilia. To answer this question, we expressed ARL3::HA::YFP in *arl3* cells at WT ARL3 level of CC-5325 control cells (resulting strain *arl3*::ARL3::HA::YFP) (Fig. S3 D). The *arl3*::ARL3::HA::YFP cells retained ARL3::HA::YFP in cilia at WT ARL3 level (Fig. 1 E) and ARL3::HA::YFP diffused into cilia to reside along the whole length of cilia as reflected by TIRF assay (Fig. 1 F and Movie S1).

**Figure 1.**
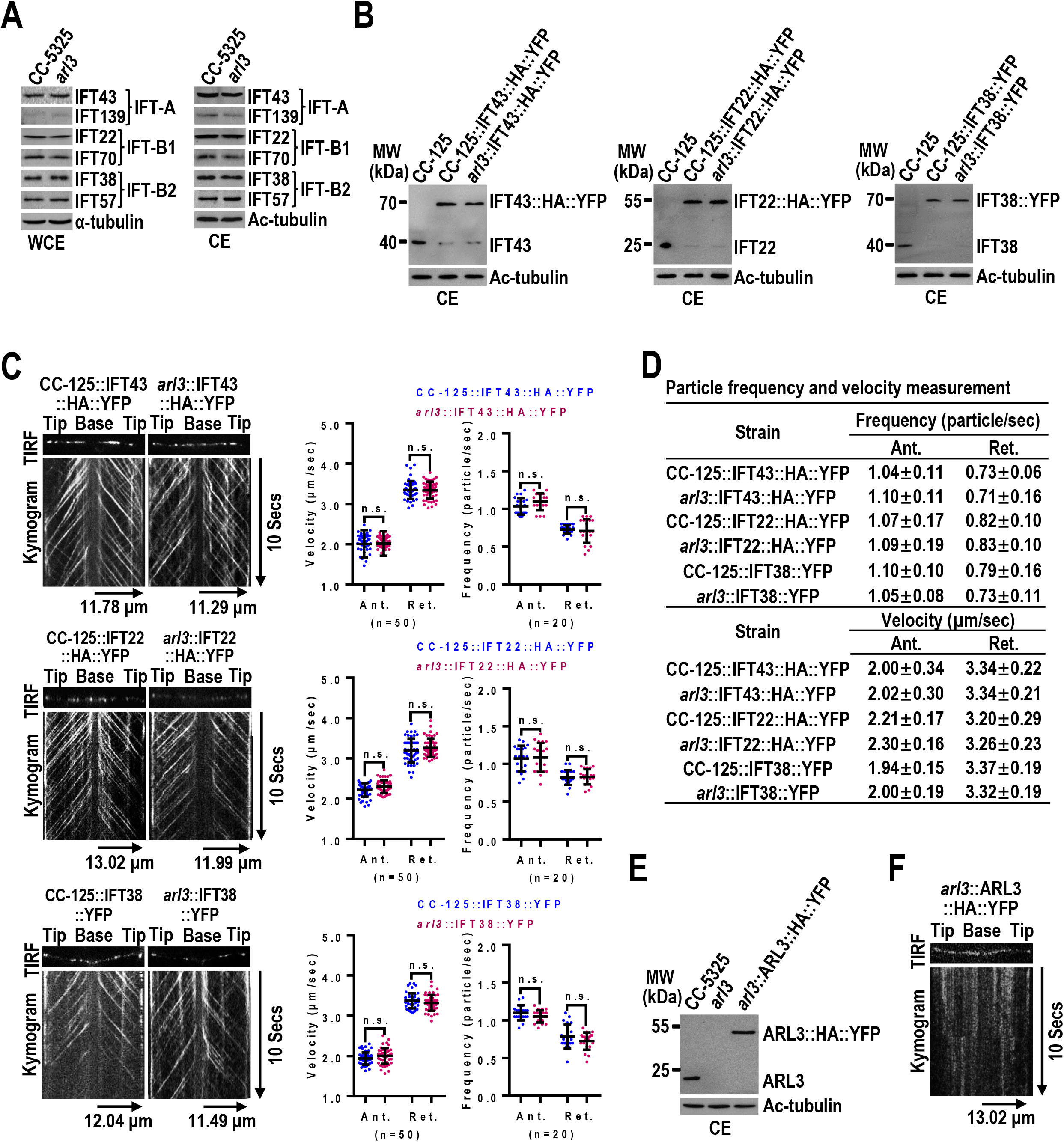
ARL3 diffuses into cilia. **(A)**. Immunoblots of whole cell extracts (WCE) and ciliary extracts (CE) of CC-5325 and *arl3* cells probed for IFT43 and IFT139 (IFT-A), IFT22 and IFT70 (IFT-B1), and IFT38 and IFT57 (IFT-B2). (**B)**. Immunoblots of CE of three cell groups including CC-125, CC-125::IFT43::HA::YFP, and *arl3*::IFT43::HA::YFP cells; CC-125, CC-125::IFT22::HA::YFP, and *arl3*::IFT22::HA::YFP cells; and CC-125, CC-125::IFT38::YFP, and *arl3*::IFT38::YFP cells probed with α-IFT43, α-IFT22, and α-IFT38, respectively. **(C)**. TIRF images and corresponding kymograms of three cell groups including CC-125::IFT43::HA::YFP and *arl3*::IFT43::HA::YFP cells; CC-125::IFT22::HA::YFP and *arl3*::IFT22::HA::YFP cells; and CC-125::IFT38::YFP and *arl3*::IFT38::YFP cells. The time and transport lengths are indicated on the right and on the bottom, respectively. The ciliary base (base) and tip (tip) were shown. Velocities and frequencies of YFP- and HA::YFP-labeled proteins to traffic inside cilia were shown as graphs. Error bar indicates S.D. n: number of cilia analyzed. n.s.: non-significance. **(D)**. Velocities and frequencies of YFP- and HA::YFP-tagged proteins to traffic inside cilia in panel C were shown as numbers. **(E)**. Immunoblots of CE of CC-5325, *arl3*, and *arl3*::ARL3::HA::YFP cells probed with α-ARL3. **(F)**. TIRF image and corresponding kymogram of the *arl3*::ARL3::HA::YFP cell (Movie S1, 15 fps). For panels **A**, **B**, and **E**, α-tubulin and acetylated (Ac)-tubulin were used to adjust the loading of WCE and CE, respectively. For panels **B** and **E**, MW: molecular weight. For panels **C** and **D**, Ant. and Ret. represent anterograde and retrograde, respectively.

### ARL3^GDP^ requires its N-terminal 15 amino acids and the G2 residue for membrane association and diffusion into cilia

Arf family GTPases associate with the inner membrane via its N-terminal amphipathic helix (Amor et al., 1994; Liu et al., 2010; Zhang et al., 2011b) and their N-terminal 15 residues are found essential for this association (Jin et al., 2010; Liu et al., 2021; Mourão et al., 2014). They were further reported to anchor to the membrane through myristylation on their glycine residue at the second amino acid position and disruption of myristylation by introducing a G2A mutation fully prevents them from associating with the membrane (Fig. S1 A) (Sahin et al., 2008; Vaughan and Moss, 1997). In addition, studies have identified the GTP-bound configuration as a prerequisite for Arf GTPases, i.e., ARL6/BBS3, to bind the membrane (Liu et al., 2021; Mourão et al., 2014). To dissect whether and how N-terminal residues, the G2 residue, and the nucleotide state confer ARL3 to bind the membrane, we applied bacteria to express ARL3 and a total of eight ARL3 variants, which contain the N-terminal 15 residue deletion (△N15), the G2A mutation, or neither combined with the mutation Q70L, T30N, or none of them (Fig. 2 A). Q70L and T30N were introduced in ARL3 or its variants as they are constitutive-active (Q70L) and dominant-negative (T30N) mutations that can lock ARL3 in the GTP- and GDP-bound state, respectively (Veltel et al., 2008). When incubated with the synthetic liposomes, ARL3 associated with liposomes only in the presence of GDP, which locks ARL3 in a GDP-bound state (Fig. 2 B). Consistent with this observation, ARL3^T30N^ rather than ARL3^Q70L^ bound liposomes, revealing that ARL3, unlike ARL6/BBS3 that binds the membrane in a GTP-dependent manner, instead relies on GDP for membrane association (Fig. 2 B) (Liu et al., 2021; Veltel et al., 2008). In contrast, ARL3△N15 and ARL3^G2A^ both were deprived of binding liposomes and they remained unbound to liposomes even when the T30N mutation was introduced, demonstrating that the N-terminal amphipathic helix and the G2 residue both are required for ARL3 to bind the membrane (Fig. 2 B). To discern if the N-terminal 15 residues, the G2 residue, and GDP are essential for ARL3 to enter cilia, we expressed the above-mentioned ARL3 variants fused at their C-terminus to HA::YFP in *arl3* cells to generate eight strains including *arl3*::ARL3^Q70L^::HA::YFP, *arl3*::ARL3^T30N^::HA::YFP, *arl3*::ARL3ΔN15::HA::YFP, *arl3*::ARL3ΔN15^Q70L^::HA::YFP, *arl3*::ARL3ΔN15^T30N^::HA::YFP, *arl3*::ARL3^G2A^::HA::YFP, *arl3*::ARL3^G2AQ70L^::HA::YFP, and *arl3*::ARL3^G2AT30N^::HA::YFP (Fig. 2 C). When expressed at WT ARL3 levels of CC-5325 control cells, ARL3^Q70L^::HA::YFP and ARL3^T30N^::HA::YFP both, unexpectedly, resembled ARL3::HA::YFP to enter cilia (Fig. 2 D). In contrast, depletion of N-terminal 15 residues and the G2A mutation alone prevented ARL3::HA::YFP from entering cilia, while Q70L rather than T30N mutation enabled both ARL3ΔN15::HA::YFP and ARL3^G2A^::HA::YFP to enter cilia (Fig. 2 D). Upon entering cilia, ARL3^T30N^::HA::YFP existed in the membrane fraction, while ARL3::HA::YFP, ARL3^Q70L^::HA::YFP, ARL3ΔN15^Q70L^::HA::YFP, and ARL3^G2^AQ70^L^::HA::YFP all resided in the matrix fraction (Fig. 2 E). Unlike ARL6/BBS3 that binds and recruits the BBSome to the basal body, thus showing a BBSome-like basal body distribution pattern (Liu et al., 2021), ARL3::HA::YFP and its cilium-entering variants did not reside at the basal body as visualized by immunostaining (Fig. S4 A). TIRF assays noticed that ARL3^T30N^::HA::YFP diffused into cilia and resembled ARL3::HA::YFP to reside along the whole length of cilia (Fig. 1 F, 2 F, and Movies S1 and S2). In contrast, ARL3^Q70L^::HA::YFP, ARL3ΔN15^Q70L^::HA::YFP, and ARL3^G2AQ70L^::HA::YFP diffused into cilia, while they mostly resided at a proximal ciliary region likely above the basal bodies (Fig. 2 F and Movies S3-S5). We failed to visualize their co-localization with the CEP290-labeled TZ region by immunostaining probably owing to their low ciliary abundance (Fig. 2 G), while we combined all these data together to conclude that ARL3 enters cilia via two different pathways. ARL3^GDP^ binds the membrane for diffusing into cilia, while ARL3^GTP^ diffuses into cilia independent of membrane association. Considering that ARL3::HA::YFP has a ciliary distribution pattern similar to its GDP-locked counterpart (Fig. 1 F and 2 F), ARL3, under physiological conditions, likely exists in a GDP-bound state and thus diffuses into cilia via the membrane association pathway in *C. reinhardtii*. Upon inside cilia, ARL3^GDP^ was quickly converted to ARL3^GTP^ by an unknown mechanism. That is why we observed ARL3::HA::YFP to reside along the whole length of cilia as shown by living TIRF assay, while it instead exists in the matrix but not membrane fraction of cilia in the steady state following ciliary fraction isolation (Fig. 2 E).

**Figure 2.**
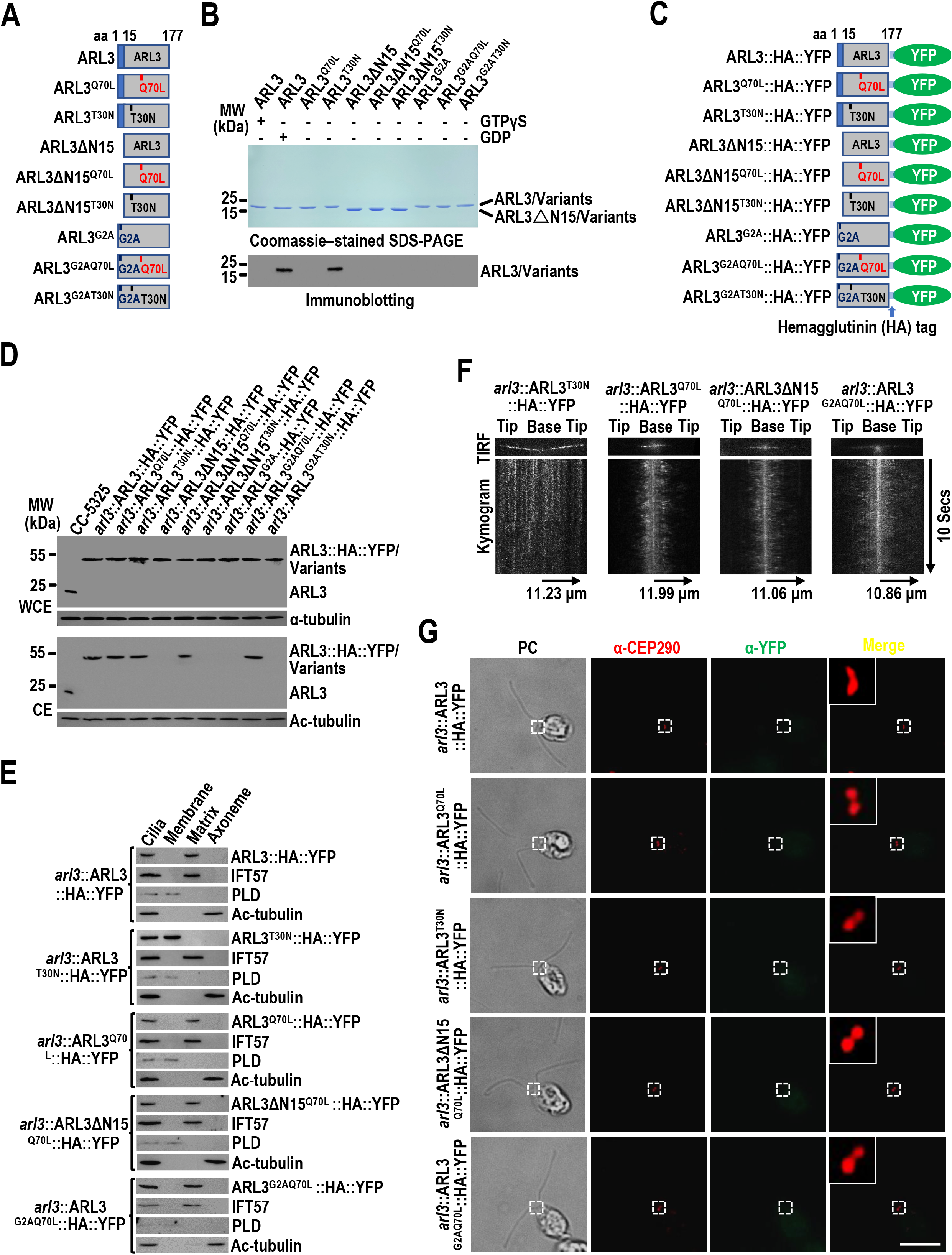
ARL3^GDP^ requires its N-terminal 15 amino acids and G2 residue for membrane association and diffusion into cilia. **(A)**. Schematic presentation of bacterially expressed ARL3 and its variants. **(B)**. SDS-PAGE visualization of bacterially expressed ARL3 and its variants indicated on the top and immunoblots of liposome-incubated ARL3 and its variants in the presence of GTPγS or GDP probed with α-ARL3. MW: molecular weight. **(C)**. Schematic presentation of ARL3::HA::YFP and its variants. **(D)**. Immunoblots of WCE and CE of cells indicated on the top probed with α-ARL3. Alpha-tubulin and acetylated (Ac)-tubulin were used to adjust the loading of WCE and CE, respectively. MW: molecular weight. **(E)**. Immunoblots of ciliary fractions of cells indicated on the left probed with α-HA, α-IFT57 (ciliary matrix marker), α-PLD (ciliary membrane marker) and Ac-tubulin (axoneme marker). **(F)**. TIRF images and corresponding kymograms of cells indicated on the top (Movies S2-S5, 15 fps). The time and transport lengths are indicated on the right and on the bottom, respectively. The ciliary base (base) and tip (tip) were shown. **(G)**. Cells indicated on the left stained with α-CEP290 (red) and α-YFP (green). Phase contrast (PC) images of cells were shown. Inset shows the proximal ciliary region. Scale bars: 10 µm. For panels **A** and **C**, △N15 stands for the N-terminal 15 amino acids of ARL3 deleted.

### ARL3^GTP^ is required for the BBSome to move cross the TZ for ciliary retrieval

Previous studies have identified ARL3 to be essential for maintaining ciliary dynamics of mouse rhodopsin and worm and human polycystin-1 and -2 (PKD1 and PKD2) (Schrick et al., 2006; Su et al., 2014; Zhang et al., 2013). These transmembrane signaling proteins cycle through cilia by IFT through binding the BBSome directly (Abd-El-Barr et al., 2007; Liu et al., 2020; Nachury, 2018; Nishimura et al., 2004; Su et al., 2014). Given that *Chlamydomonas* ARL3 does not affect IFT, we wondered whether ARL3 could mediate signaling protein dynamics in cilia via the BBSome pathway. To answer this question, we examined *arl3* cells and the rescuing strains *arl3*::ARL3::HA::YFP, *arl3*::ARL3^Q70L^::HA::YFP, and *arl3*::ARL3^T30N^::HA::YFP. As compared to CC-5325 control cells, four strains retained the BBSome subunits BBS1, BBS4, BBS5, BBS7, and BBS8 at WT levels (Fig. 3 A). Of note, these BBSome subunits accumulated in *arl3* cilia to levels ∼7.5-fold higher than control cell cilia (Fig. 3 B). This observation was confirmed as rescuing ARL3 with ARL3::HA::YFP restored them back to normal, as shown in *arl3*::ARL3::HA::YFP cilia (Fig. 3 B). We further identified ARL3^Q70L^::HA::YFP rather than ARL3^T30N^::HA::YFP is able to restore these BBSome subunits back to normal in *arl3* cilia, revealing that ARL3^GTP^ is required for maintaining BBSome ciliary dynamics (Fig. 3 B). Our previous study has shown that the BBSome disassembles at the ciliary tip followed by a reassembly process for loading onto retrograde IFT trains for transporting to the ciliary base (Sun et al., 2021). In the absence of ARL3, the BBSome remained as an intact entity in cilia, excluding ARL3 from mediating BBSome remodeling at the ciliary tip (Fig. 3 C). To further discern how ARL3 mediates BBSome dynamics in cilia, we generated a ARL3- and BBS8-double null mutant that we named *arl3-bbs8* and expressed BBS8::YFP in *arl3-bbs8* cells (resulting strain *arl3-bbs8*::BBS8::YFP) at BBS8::YFP level of *bbs8*::BBS8::YFP cells, which expresses BBS8::YFP in BBS8-null *bbs8* cells (Fig. 3 D). When expressed at WT BBS8 level of CC-125 control cells, BBS8::YFP entered and retained in *bbs8*::BBS8::YFP cilia at the endogenous BBS8 level of control cells and so did for the BBSome subunits BBS1, BBS4, BBS5, and BBS7 (Fig. 3 D). In contrast, ARL3 knockout did not affect cellular levels of BBS1, BBS4, BBS5, and BBS7 but caused them and BBS8::YFP to build up in *arl3-bbs8*::BBS8::YFP cilia (Fig. 3 D). Given that the BBSome (represented by BBS8::YFP) underwent typical bidirectional IFT of *Chlamydomonas* BBSome the same in cilia of both transgenic cells, ARL3 was then excluded from mediating BBSome transportation between the ciliary tip and base (Fig. 3 E and F and Movies S6-S7). Interestingly, BBS8::YFP was instead visualized to accumulate right above the CEP290-labled TZ region of *arl3-bbs8*::BBS8::YFP cilia as compared to *bbs8*::BBS8::YFP cilia (Fig. 3 G). This buildup was defined to a proximal ciliary region obviously above the IFT46-labled basal bodies (Fig. S4 B). Taking these data together, we conclude that ARL3 is dispensable for the BBSome to enter and traffic inside cilia, while the BBSome detaches from retrograde IFT at the proximal ciliary region right above the TZ, and it requires ARL3^GTP^ for moving cross the TZ and out of cilia. This notion was verified as the endogenous BBS8 in *arl3* and *arl3*::ARL3^T30N^::HA::YFP cilia accumulated at the proximal ciliary region obviously above the IFT81-labled basal bodies as compared to CC-5325, *arl3*::ARL3::HA::YFP, and *arl3*::ARL3^Q70L^::HA::YFP cilia (Fig. S4 C).

**Figure 3.**
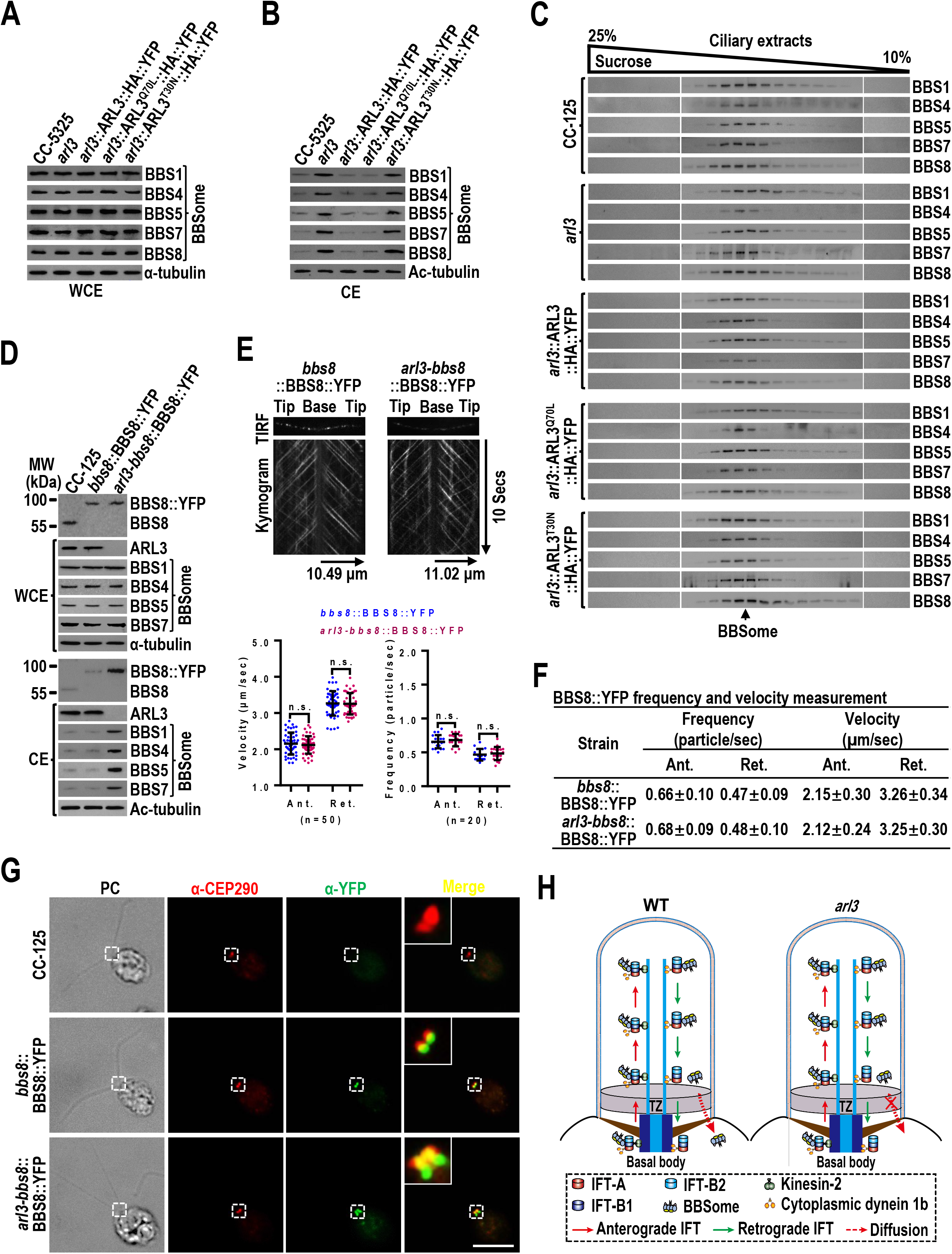
ARL3^GTP^ is required for the BBSome to move cross the TZ for ciliary retrieval. **(A and B)**. Immunoblots of WCE (**A**) and CE (**B**) of cells indicated on the top probed for the BBSome subunits BBS1, BBS4, BBS5, BBS7, and BBS8. **(C)**. Immunoblots of sucrose density gradient of CE of cells indicated on the left probed for the BBSome subunits BBS1, BBS4, BBS5, BBS7, and BBS8. **(D)**. Immunoblots of WCE and CE of cells indicated on the top probed with α-BBS8, α-ARL3, α-BBS1, α-BBS4, α-BBS5, and α-BBS7. MW: molecular weight. **(E)**. TIRF images and corresponding kymograms of *bbs8*::BBS8::YFP and *arl3-bbs8*::BBS8::YFP cells (Movies S6-S7, 15 fps). The time and transport lengths are indicated on the right and on the bottom, respectively. The ciliary base (base) and tip (tip) were shown. Velocities and frequencies of fluorescent proteins to traffic inside cilia were shown as graphs. Error bar indicates S.D. n: number of cilia analyzed. n.s.: non-significance. **(F)**. Velocities and frequencies of BBS8::YFP to traffic inside cilia of *bbs8*::BBS8::YFP and *arl3-bbs8*::BBS8::YFP cells in panel **E** were shown as numbers. **(G)**. CC-125, *bbs8*::BBS8::YFP, and *arl3-bbs8*::BBS8::YFP cells stained with α-CEP290 (red) and α-YFP (green). Phase contrast (PC) images of cells are shown. Inset shows the proximal ciliary region. Scale bars: 10 µm. H. Schematic representation of how loss of ARL3 blocks outward diffusion of the BBSome through the TZ for ciliary retrieval. For panels **A**, **B**, and **D**, alpha-tubulin and acetylated (Ac)-tubulin were used to adjust the loading of WCE and CE, respectively. For panels **E** and **F**, Ant. and Ret. represent anterograde and retrograde, respectively.

**Figure 4.**
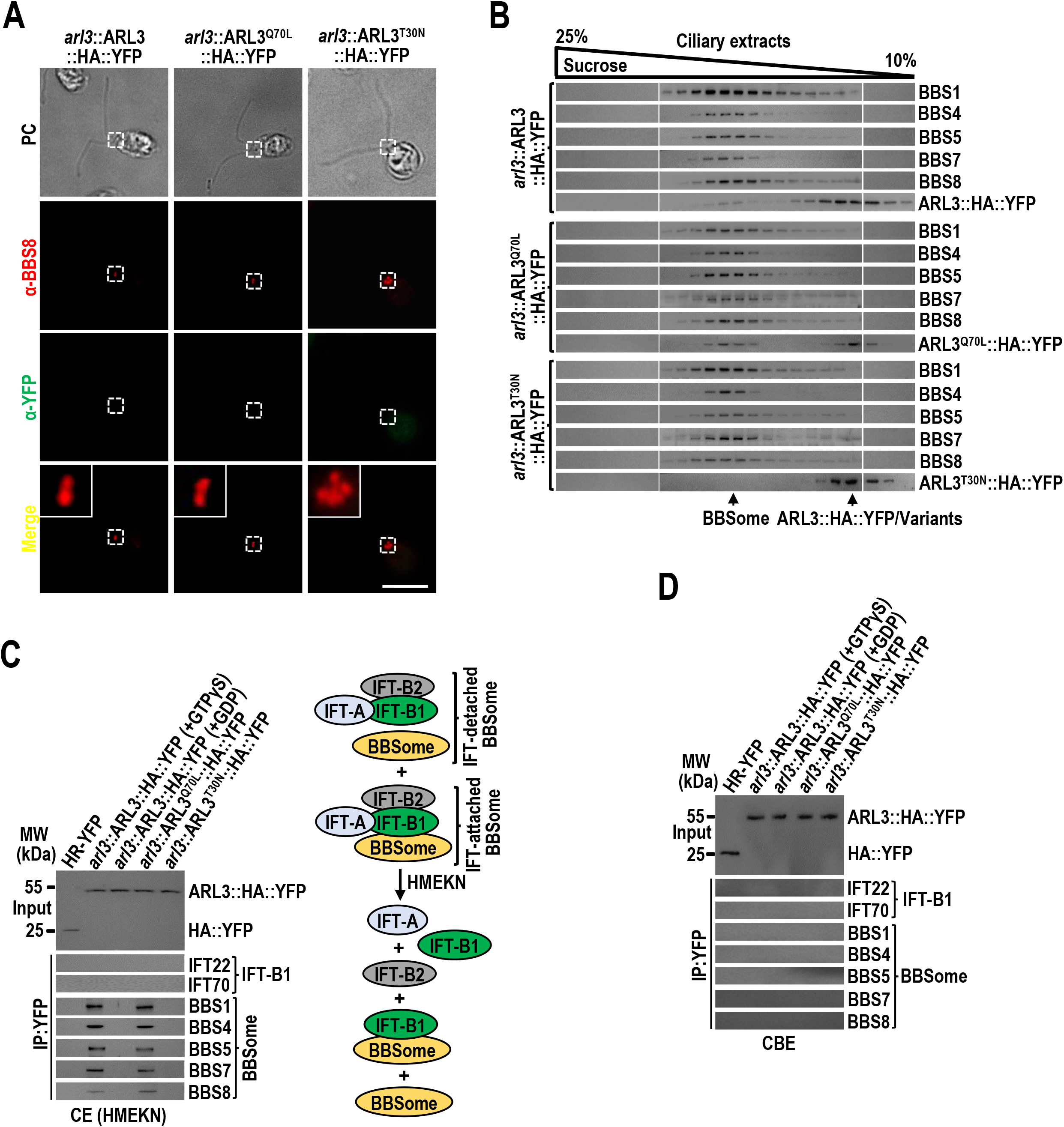
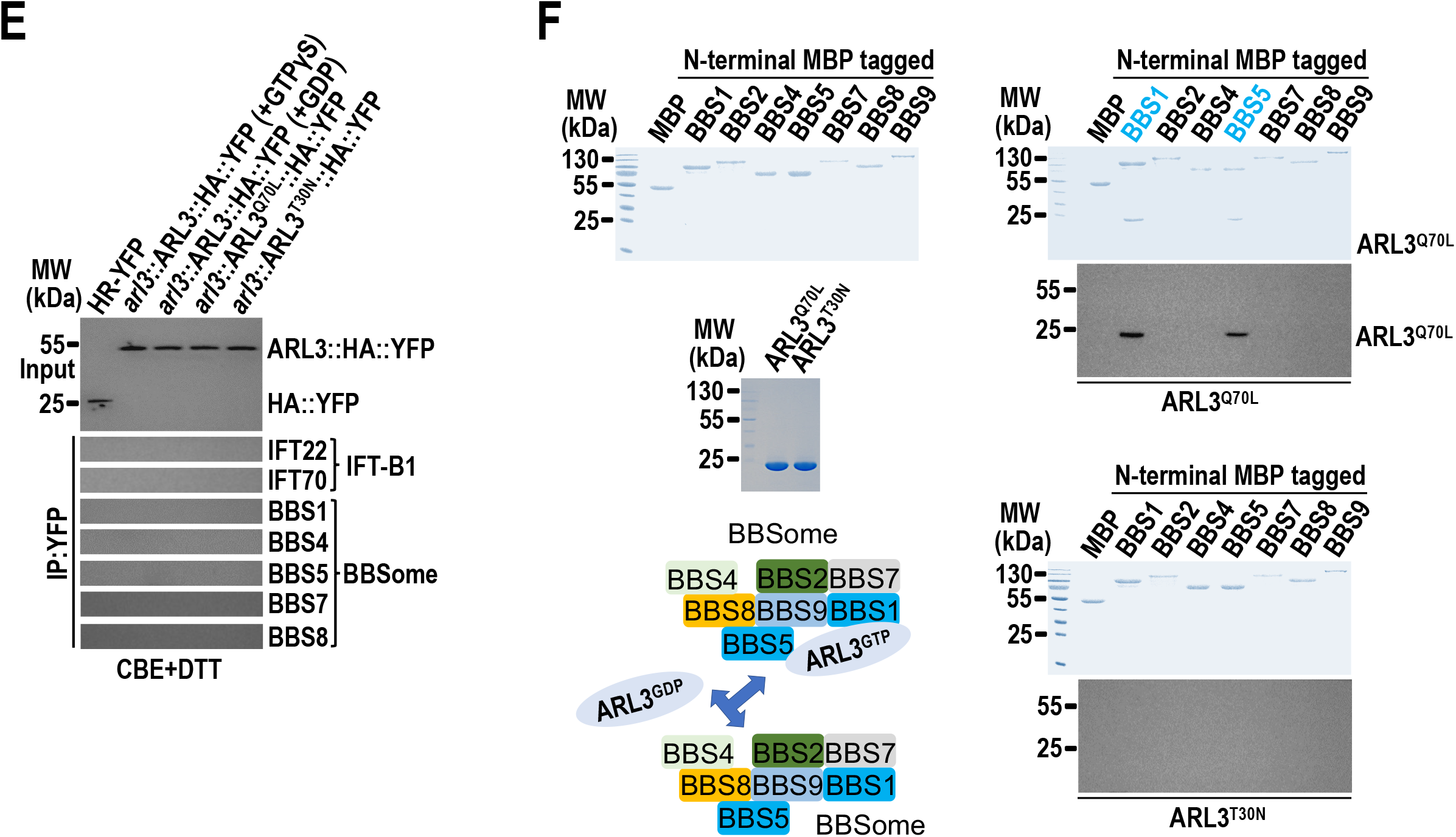
The BBSome is a major ARL3 effector at the proximal ciliary region but not in the cell body. **(A)**. *arl3*::ARL3::HA::YFP, *arl3*::ARL3^Q70L^::HA::YFP, and *arl3*::ARL3^T30N^::HA::YFP cells stained with α-BBS8 (red) and α-YFP (green). Phase contrast (PC) images of cells were also shown. Inset shows the proximal ciliary region and the basal bodies. Scale bars: 10 µm. **(B)**. Immunoblots of sucrose density gradient of CE of *arl3*::ARL3::HA::YFP, *arl3*::ARL3^Q70L^::HA::YFP, and *arl3*::ARL3^T30N^::HA::YFP cells probed with α-BBS1, α-BBS4, α-BBS5, α-BBS7, α-BBS8, and α-ARL3. **(C)**. Immunoblots of α-YFP-captured proteins from CE of HR-YFP (HA::YFP-expressing CC-125 cells), *arl3*::ARL3::HA::YFP (in the presence of GTPγS or GDP), *arl3*::ARL3^Q70L^::HA::YFP, and *arl3*::ARL3^T30N^::HA::YFP cells probed for the IFT-B1 subunits IFT22 and IFT70 and the BBSome subunits BBS1, BBS4, BBS5, BBS7, and BBS8. Input was quantified with α-YFP by immunoblotting. A schematic representation of how IFT-detached BBSome exists independently of the IFT-B1/BBSome entity in HMEKN buffer was shown on the right. **(D and E)**. Immunoblots of α-YFP-captured proteins from cell body extracts (CBE) of HR-YFP (HA::YFP-expressing CC-125 cells), *arl3*::ARL3::HA::YFP (in the presence of GTPγS or GDP), *arl3*::ARL3^Q70L^::HA::YFP, and *arl3*::ARL3^T30N^::HA::YFP cells probed for the IFT-B1 subunits IFT22 and IFT70 and the BBSome subunits BBS1, BBS4, BBS5, BBS7, and BBS8 in the absence of DTT (**D**) and in the presence of DTT (**E**). For both panels, input was quantified with α-YFP by immunoblotting. MW stands for molecular weight. **(F)**. Bacterially expressed MBP, MBP::BBS1, MBP::BBS2, MBP::BBS4, MBP::BBS5, MBP::BBS7, MBP::BBS8, and MBP::BBS9 (upper left) were mixed with ARL3^Q70L^ or ARL3^T30N^ (middle left) and complexes recovered on amylose beads were resolved by SDS-PAGE followed by Coomassie blue staining and immunoblotting with α-ARL3 (right). A schematic representation of direct interactions of ARL3^Q70L^ but not ARL3^T30N^ with BBS1 and BBS5 of the BBSome was shown (lower left). MW, molecular weight.

### The BBSome is a major ARL3 effector at the proximal ciliary region but not in the cell body

Different from ARL3^GDP^ that binds the ciliary membrane, ARL3^GTP^ resembles the BBSome to reside in the ciliary matrix (Fig. 2 E). ARL3^GDP^ resides along the whole length of cilia, while GTP loading largely restricts ARL3 to the proximal ciliary region, where the BBSome sheds from IFT and accumulates in the absence of ARL3 (Fig. 3 G and Fig. S4 B and C). As expected, neither ARL3::HA::YFP nor the two mutants can be visualized by immunostaining, preventing us from drawing a conclusion that they co-localize with the BBSome (represented by BBS8) at the proximal ciliary region right above the TZ (Fig. 4 A), while the observation that ARL3^GTP^ promotes outward movement of the BBSome across the TZ proposed that ARL3^GTP^ may achieve this goal via interacting with the BBSome (Fig. 3). This notion was supported as partial ARL3::HA::YFP and ARL3^Q70L^::HA::YFP co-sedimented with the BBSome in *arl3*::ARL3::HA::YFP and *arl3*::ARL3^Q70L^::HA::YFP cilia in sucrose density gradients (Fig. 4 B). In contrast, ARL3^T30N^::HA::YFP remained to be separated from the BBSome in *arl3*::ARL3^T30N^::HA::YFP cilia, as shown by sucrose density gradient centrifugation assays (Fig. 4 B). Our previous studies have shown that HMEKN buffer confers IFT-A, IFT-B1, and IFT-B2 subcomplexes to separate from one another, while the BBSome remains to be associated with IFT-B1 (Sun et al., 2021). In the presence of GTPγS that locks ARL3 in a GTP-bound state, ARL3::HA::YFP immunoprecipitated the BBSome subunits BBS1, BBS4, BBS5, BBS7, and BBS8 but not the IFT-B1 subunits IFT22 and IFT70 in *arl3*::ARL3::HA::YFP cilia (Fig. 4 C). In contrast, none of these proteins were recovered in the presence of GDP that locks ARL3 in a GDP-bound state, identifying IFT-B1-separated BBSomes indeed exist in cilia for ARL3^GTP^ to interact with (Fig. 4 C). This notion was verified as ARL3^Q70L^::HA::YFP but not ARL3^T30N^::HA::YFP recovered the BBSome and none of them was able to recover IFT-B1, as shown in *arl3*::ARL3^Q70L^::HA::YFP and *arl3*::ARL3^T30N^::HA::YFP cilia (Fig. 4 C). Other than this, we further identified ARL3^Q70L^::HA::YFP fails to immunoprecipitate the BBSome subunits nor IFT proteins in the cell body extracts even in the presence of dithiothreitol (DTT) that separates the BBSome from IFT-B1 B1 (Sun et al., 2021), revealing that ARL3^GTP^ interaction with the IFT-detached BBSome only occurs in cilia but not in the cell body (Fig. 4 D and E). As identified by in vitro protein interaction assays, BBS1 and BBS5 were shown to be the BBSome subunits most efficiently captured by ARL3^Q70L^ but not ARL3^T30N^ *in vitro*, revealing that ARL3, only when in a GTP-bound state, interacts with the IFT-separated BBSome directly (Fig. 4 F). We then conclude that the BBSome is the major ARL3 effector only when they both position at the proximal ciliary region.

### ARL3^GTP^ recruits the BBSome for diffusing through the TZ for ciliary retrieval

ARL3^GTP^ promotes outward BBSome movement cross the TZ (Fig. 3). To have a full review on how ARL3 and the BBSome interplay for ciliary retrieval, we ought to dissect whether ARL3 ciliary dynamics is mediated by the BBSome. To solve this puzzle, we expressed ARL3::HA::YFP, ARL3^Q70L^::HA::YFP, and ARL3^T30N^::HA::YFP in ARL3-and BBS8-double null *arl3-bbs8* cells to generate three strains *arl3-bbs8*::ARL3::HA::YFP, *arl3-bbs8*::ARL3^Q70L^::HA::YFP, and *arl3-bbs8*::ARL3^T30N^::HA::YFP. By doing so, the BBSome was deprived of ciliary presence in these cells as BBS8 knockout disrupts BBSome assembly in the cell body, thus making the BBSome unavailable for entering cilia (Fig. 5 A). When expressed at WT ARL3 level of CC-125 control cells, ARL3::HA::YFP, ARL3^Q70L^::HA::YFP, and ARL3^T30N^::HA::YFP entered and retained at WT ARL3 level in cilia (Fig. 5 A). Like in ARL3-null *arl3* cells, ARL3^T30N^::HA::YFP resided in the membrane fraction, while ARL3::HA::YFP and ARL3^Q70L^::HA::YFP both existed in the matrix fraction in the absence of the BBSome (Fig. 5 B). Immunostaining, as expected, was unable to visualize these recombinant proteins to reside at the CEP290-labelled TZ region (Fig. S4 D). However, TIRF assays visualized that, like in *arl3* cells, all three fluorescent proteins entered cilia by diffusion in the *arl3-bbs8* double mutant cells (Fig. 5 C). Once inside cilia, ARL3::HA::YFP and ARL3^T30N^::HA::YFP distributed to the whole length of cilia, while ARL3^Q70L^::HA::YFP mostly concentrated at the proximal ciliary region (Fig. 5 C). These data thus excluded the BBSome from mediating ARL3 ciliary dynamics.

**Figure 5.**
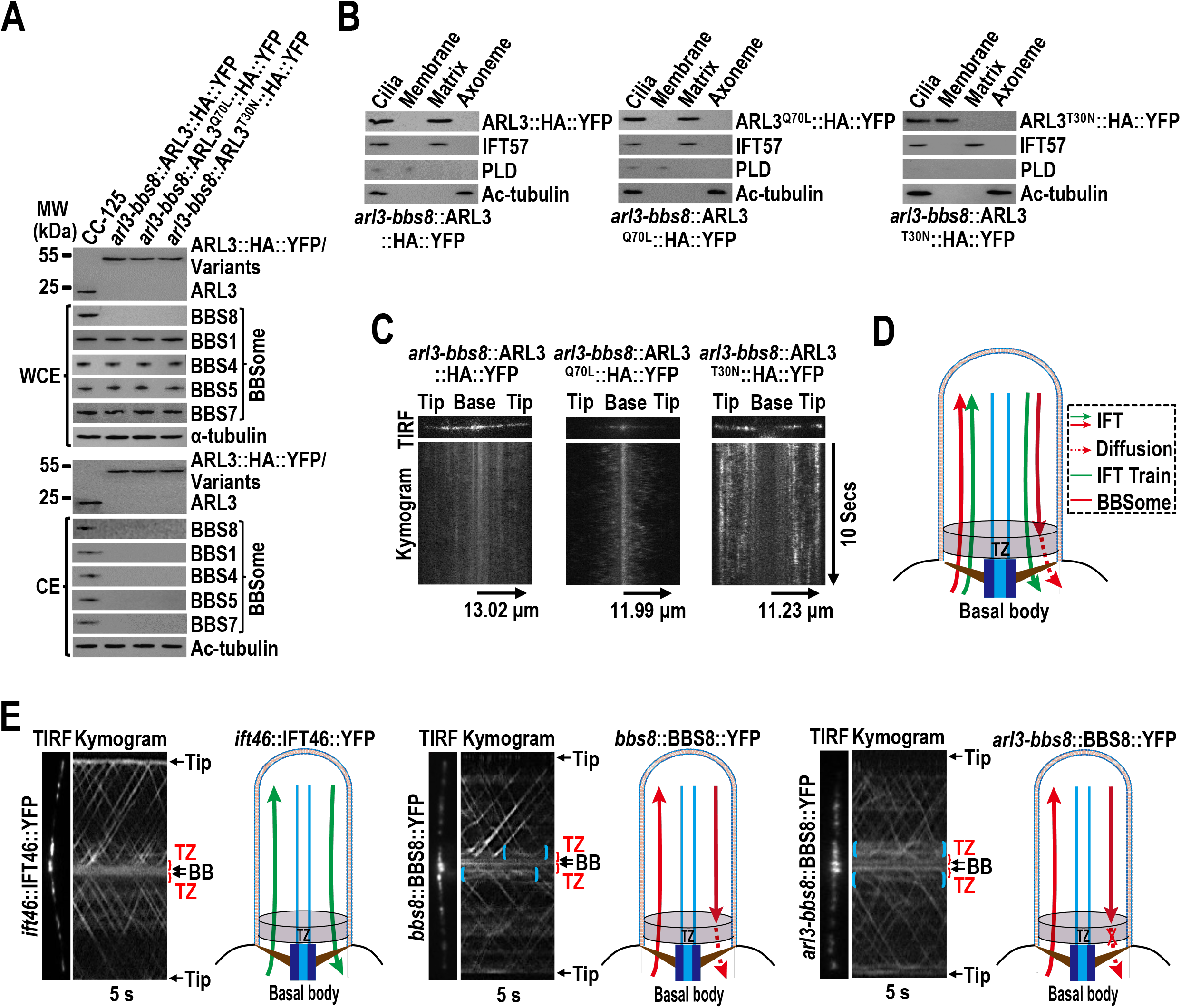
ARL3^GTP^ recruits the BBSome for diffusing through the TZ for ciliary retrieval. **(A)**. Immunoblots of WCE and CE of cells indicated on the top probed with α-ARL3, α-BBS8, α-BBS1, α-BBS4, α-BBS5, and α-BBS7. Alpha-tubulin and acetylated (Ac)-tubulin were used to adjust the loading of WCE and CE, respectively. MW: molecular weight. **(B)**. Immunoblots of ciliary fractions of *arl3-bbs8*::ARL3::HA::YFP, *arl3-bbs8*::ARL3^Q70L^::HA::YFP, and *arl3-bbs8*::ARL3^T30N^::HA::YFP cells probed with α-HA, α-IFT57 (ciliary matrix marker), α-PLD (ciliary membrane marker) and Ac-tubulin (axoneme marker). **(C)**. TIRF images and corresponding kymograms of *arl3-bbs8*::ARL3::HA::YFP, *arl3-bbs8*::ARL3^Q70L^::HA::YFP, and *arl3-bbs8*::ARL3^T30N^::HA::YFP cells. The time and transport lengths are indicated on the right and on the bottom, respectively. The ciliary base (base) and tip (tip) were shown. **(D)**. Schematic representation of how IFT trains and the BBSome cycle between basal body and cilia. BBSome diffusion through the TZ for ciliary retrieval was shown. **(E)**. TIRF images and corresponding kymograms of *ift46*::IFT46::YFP, *bbs8*::BBS8::YFP, and *arl3-bbs8*::BBS8::YFP cells (Movies S8-S10, 15 fps). The time was indicated on the bottom. The ciliary base (base) and tip (tip), the transition zone (TZ) and the basal body (BB) were shown. The corresponding schematic representation of how IFT46::YFP and BBS8::YFP cycle between the basal body and cilia was shown.

Knockout of *Chlamydomonas* ARL3 does not disrupt IFT but causes BBSome accumulation at the proximal ciliary region right above the TZ, consistent with the observation that the human and murine BBSome sheds from retrograde IFT before diffusing through the TZ for ciliary retrieval (Fig. 5 D) (Nachury, 2018; Ye et al., 2018). To visualize whether ARL3^GTP^ promotes BBSome diffusion through the TZ for ciliary retrieval in *C. reinhardtii*, we examined the IFT46::YFP-expressing *ift46*::IFT46::YFP and the BBS8::YFP-expressing *bbs8*::BBS8::YFP and *arl3-bbs8*::BBS8::YFP cells (Lv et al., 2017). Of note, TIRF assays identified retrograde IFT trains (represented by IFT46::YFP) transported from the ciliary tip all the way to the basal bodies, suggesting that they move cross the TZ for ciliary retrieval via IFT (Fig. 5 E and Movie S8). The BBSome (represented by BBS8::YFP) performed normal IFT for trafficking from the ciliary tip to base (Fig. 5 E and Movie S9). When reaching the proximal ciliary region right above the TZ, the BBSome stopped performing IFT and shifted to diffuse through the TZ for ciliary retrieval (Fig. 5 E and Movie S9). In the absence of ARL3, the BBSome (represented by BBS8::YFP) underwent normal bidirectional IFT, while its suspension for diffusing through the TZ but accumulating at the proximal ciliary region right above the TZ was easily observed, as shown in cilia of *arl3-bbs8*::BBS8::YFP cells (Fig. 5 E and Movie S10). In summary, the BBSome relies on ARL3^GTP^ for diffusing through the TZ for ciliary retrieval but not *vice versa*.

### ARL3^GTP^ recruits PLD-laden BBSomes to move cross the TZ for ciliary retrieval

Our previous study and others have shown that the ciliary membrane anchored PLD couples with the BBSome at the ciliary tip followed by exiting cilia via IFT (Liu and Lechtreck, 2018; Liu et al., 2021). As compared to CC-125 control cells, *arl3* cells retained PLD at the WT level but accumulated it in cilia (Fig. 6 A). Immunostaining identified PLD is not able to be visualized in CC-125 cilia but accumulates to become visible at the proximal ciliary above the IFT-81-labeled basal bodies in *arl3* cells (Fig. 6 B). The PLD abundance was restored to normal in cilia of *arl3*::ARL3::HA::YFP and *arl3*::ARL3^Q70L^::HA::YFP cells but not *arl3*::ARL3^T30N^::HA::YFP cells (Fig. 6 A). Accordingly, PLD became invisible in cilia of *arl3*::ARL3::HA::YFP and *arl3*::ARL3^Q70L^::HA::YFP cells as in cilia of CC-125 control cells but remained to be accumulated at the proximal ciliary region above the IFT-81-labeled basal bodies in *arl3*::ARL3^T30N^::HA::YFP cells (Fig. 6 B). Given that ARL3^GTP^ directs the BBSome to behave the same way as PLD in these events (Fig. 3 A and B and Fig. S4 C), ARL3^GTP^ is supposed to be desirable for promoting both PLD and the BBSome to move cross the diffusion barrier at the TZ for ciliary retrieval. Notably, ARL3::HA::YFP immunoprecipitated the BBSome and PLD but not IFT-A, IFT-B1, and IFT-B2 in the presence of GTPγS that locks ARL3 in a GTP-bound state (Fig. 6 C). In the presence of GDP that locks ARL3 in a GDP-bound state, ARL3::HA::YFP recovered none of these proteins (Fig. 6 C). This observation was confirmed as ARL3^Q70L^::HA::YFP but not ARL3^30N^::HA::YFP immunoprecipitated the IFT-detached BBSome and PLD (Fig. 6 C). These data provided evidence to show that PLD remains to be a cargo of the IFT-detached BBSome during its diffusion through the TZ for ciliary retrieval (Fig. 5 E). Therefore, ARL3^GTP^ binds and recruits PLD-laden BBSomes to move cross the diffusion barrier at the TZ for ciliary retrieval.

**Figure 6.**
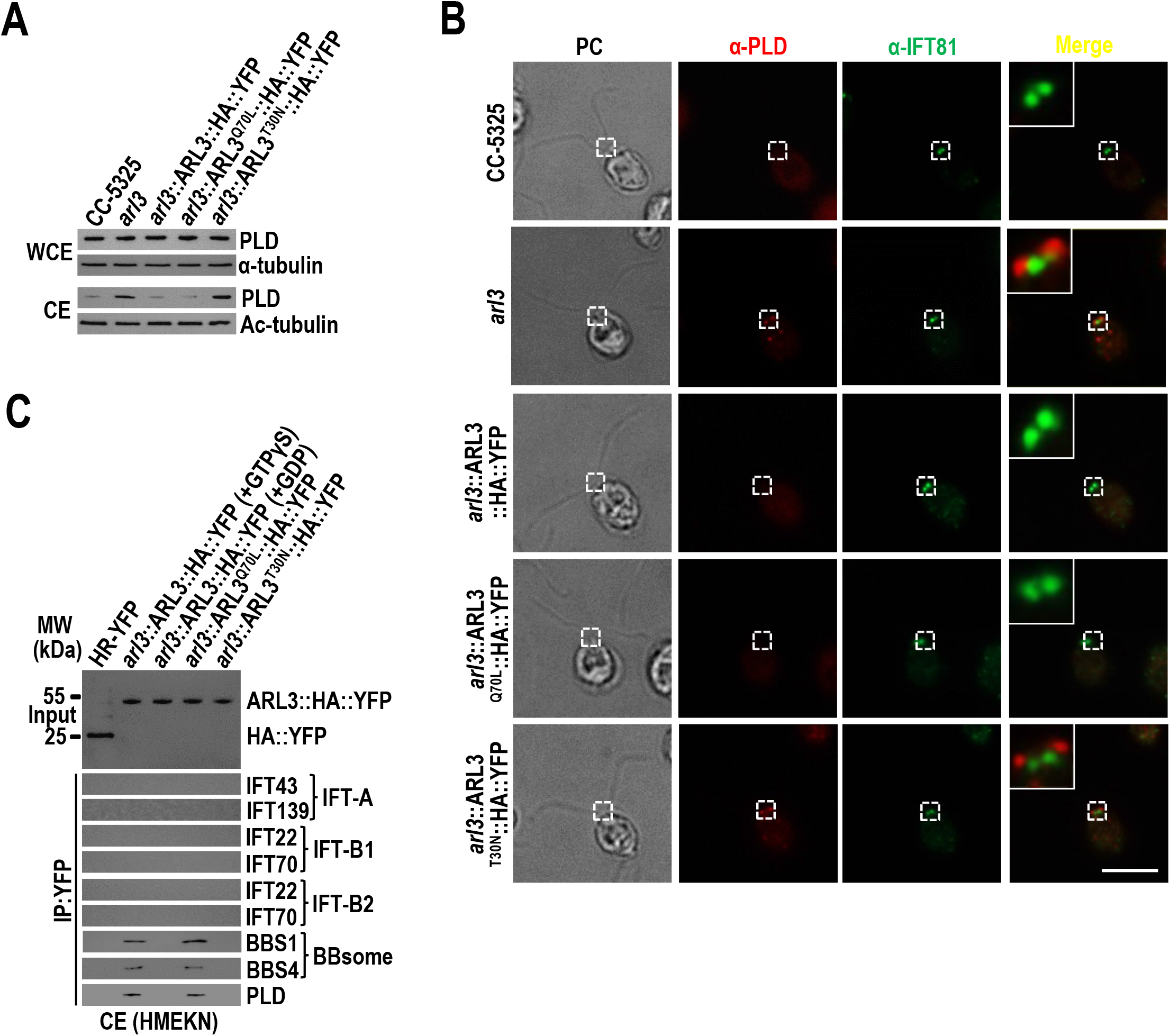
ARL3^GTP^ recruits PLD-laden BBSomes to move cross the TZ for ciliary retrieval. **(A)**. Immunoblots of WCE and CE of CC-5325, *arl3*, *arl3*::ARL3::HA::YFP, *arl3*::ARL3^Q70L^::HA::YFP, and *arl3*::ARL3^T30N^::HA::YFP cells probed for PLD. Alpha-tubulin and acetylated-α-tubulin (Ac-tubulin) were used as a loading control for WCE and CE, respectively. **(B)**. CC-5325, *arl3*, *arl3*::ARL3::HA::YFP, *arl3*::ARL3^Q70L^::HA::YFP, and *arl3*::ARL3^T30N^::HA::YFP cells were stained with α-PLD (red) and α-IFT81 (green). Phase contrast (PC) images of cells were shown. Inset shows the proximal ciliary region and the basal bodies. Scale bars: 10 µm. **(C)**. Immunoblots of α-YFP-captured proteins from CE of HR-YFP (HA::YFP-expressing CC-125 cells), *arl3*::ARL3::HA::YFP (in the presence of GTPγS or GDP), *arl3*::ARL3^Q70L^::HA::YFP, and *arl3*::ARL3::HA::YFP cells probed for the IFT-B1 subunits IFT22 and IFT70, the IFT-B2 subunits IFT38 and IFT57, the IFT-A subunits IFT43 and IFT139, the BBSome subunits BBS1 and BBS4, and PLD. Input was quantified with α-YFP by immunoblotting. MW stands for molecular weight.

### ARL3 mediates phototaxis through controlling BBSome ciliary retrieval

Our previous study and others have shown that disruption of BBSome ciliary dynamics leads to the generation of *Chlamydomonas* cells defective in phototaxis (Liu and Lechtreck, 2018; Sun et al., 2021). The *arl3* cells were disabled for conducting phototaxis as they were derived from phototaxis-deficient CC-5325 cells (Fig. 7 A and B). To examine if ARL3 mediates phototaxis through controlling BBSome ciliary dynamics, we next applied microRNA (miRNA) vector to knock down the endogenous ARL3 to ∼8.3% of WT level in CC-125 cells; we referred to this strain as ARL3^miRNA^ (Fig. 7 C). Reflecting its cellular reduction, ARL3 was strongly reduced to ∼8.0% of WT level in ARL3^miRNA^ cilia (Fig. 7 C). Consistent with ARL3 knockout result, partial depletion of ARL3 did not appear to affect ciliary length (Fig. S5 A) nor altered cellular and ciliary abundance of IFT proteins (Fig. S5 B). As expected, BBS1, BBS4, BBS5, BBS7, and BBS8 were observed to accumulate in ARL3^miRNA^ cilia, while ARL3 knockdown did not alter their cellular contents (Fig. 7 C). After the BBSome (represented by BBS8) was determined to accumulate at a proximal region of ARL3^miRNA^ cilia above the IFT81-labeled basal bodies, we concluded that ARL3 knockdown mimics ARL3 knockout for blocking BBSome to diffuse through the TZ for ciliary retrieval (Fig. 7 D). This notion was verified as rescue of ARL3 with ARL3::HA::YFP to WT level (resulting strain ARL3^Res-WT^) did not alter cellular contents of BBS1, BBS4, BBS5, BBS7, and BBS8 but restored them to WT levels in cilia (Fig. 7 C and D). We further observed that rescue of ARL3 with ARL3^Q70L^::HA::YFP but not ARL3^T30N^::HA::YFP (resulting strains ARL3^Res-Q70L^ and ARL3^Res-T30N^) restored those BBSome subunits to WT levels in cilia, verifying that GTP-loading confers ARL3 to promote the BBSome to diffuse through the TZ for ciliary retrieval (Fig. 7 C and D). After obtaining these cells, we performed both population and single cell assays to determine their phototoxic responses and identified ARL3^miRNA^ cells are non-phototactic, ARL3^Res-WT^ and ARL3^Res-Q70L^ cells, like CC-125 control cells, became normal in phototaxis, and ARL3^Res-T30N^ cells remained to be non-phototactic (Fig. 7 E and F). Therefore, ARL3 controls *Chlamydomonas* phototaxis through maintaining BBSome ciliary dynamics by controlling BBSome diffusion through the TZ for ciliary retrieval in a GTP-dependent manner.

**Figure 7.**
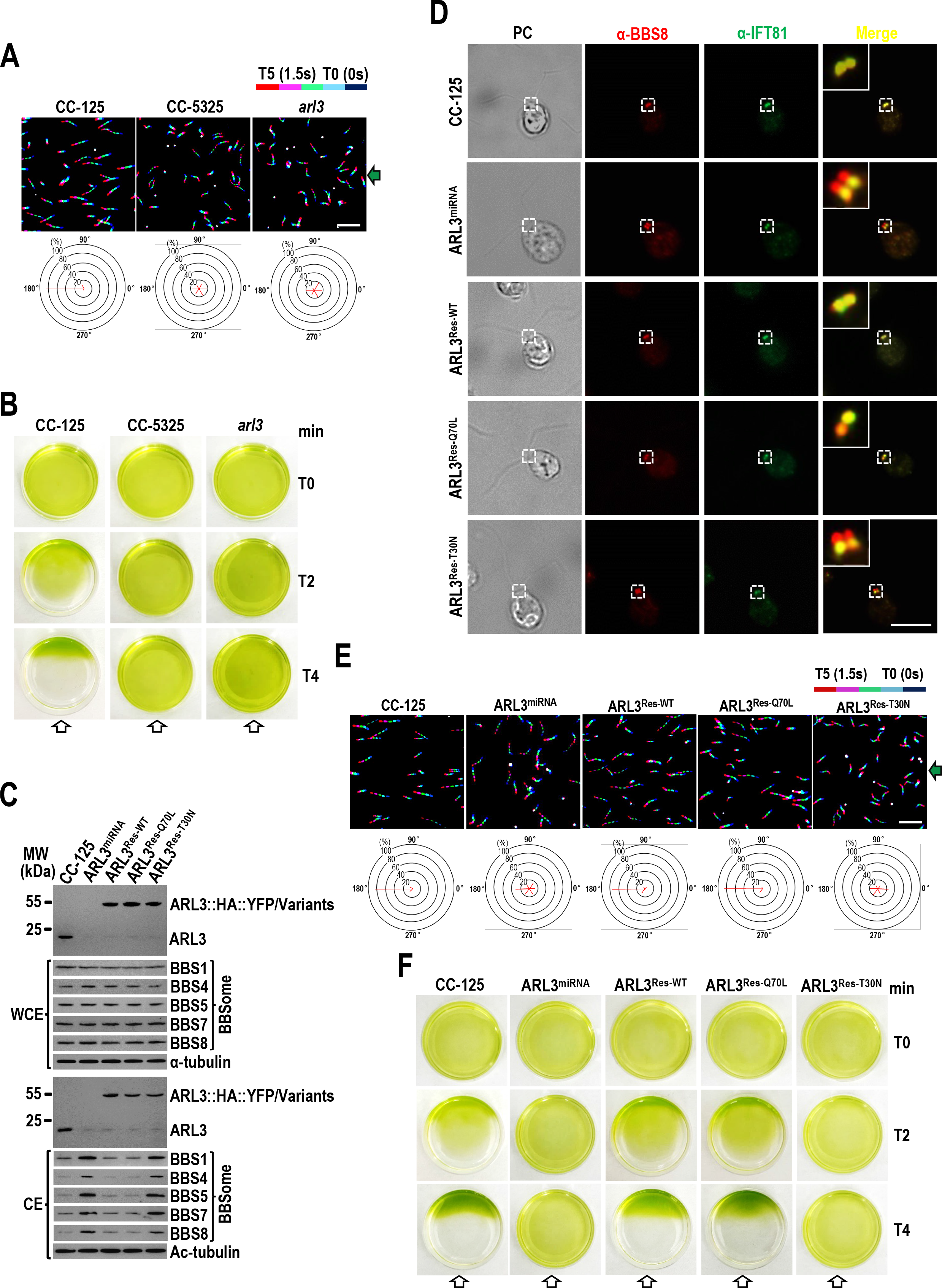
ARL3 mediates phototaxis through controlling BBSome ciliary retrieval. **(A and B)**. Single-cell motion assay (**A**) and population phototaxis assay (**B**) of CC-125, CC-5325, and *arl3* cells. **(C)**. Immunoblots of WCE and CE of cells indicated on the top probed with α-ARL3, α-BBS1, α-BBS4, α-BBS5, α-BBS7, and α-BBS8. Alpha-tubulin and acetylated (Ac)-tubulin were used to adjust the loading of WCE and CE, respectively. MW: molecular weight. **(D)**. Cells indicated on the left were stained with α-BBS8 (red) and α-IFT81 (green). Phase contrast (PC) images of cells were also shown. Inset shows the proximal ciliary region right and the basal bodies. Scale bars: 10 µm. **(E and F)**. Single-cell motion assay (**E**) and population phototaxis assay (**F**) of CC-125, ARL3^miRNA^, ARL3^Res-WT^, ARL3^Res-Q70L^, and ARL3^Res-T30N^ cells. For panels **A** and **E**, the direction of light is indicated as green arrows and the radial histograms show the percentage of cells moving in a particular direction relative to the light (six bins of 60° each). Composite micrographs show the tracks of single cells. Each of the five merged frames was assigned a different color (blue is frame 1 and red is frame 5, corresponding to a travel time of 1.5 s). Scale bar: 10 μm. For panels **B** and **F**, the direction of light is indicated as white arrows.

**Figure 8.**
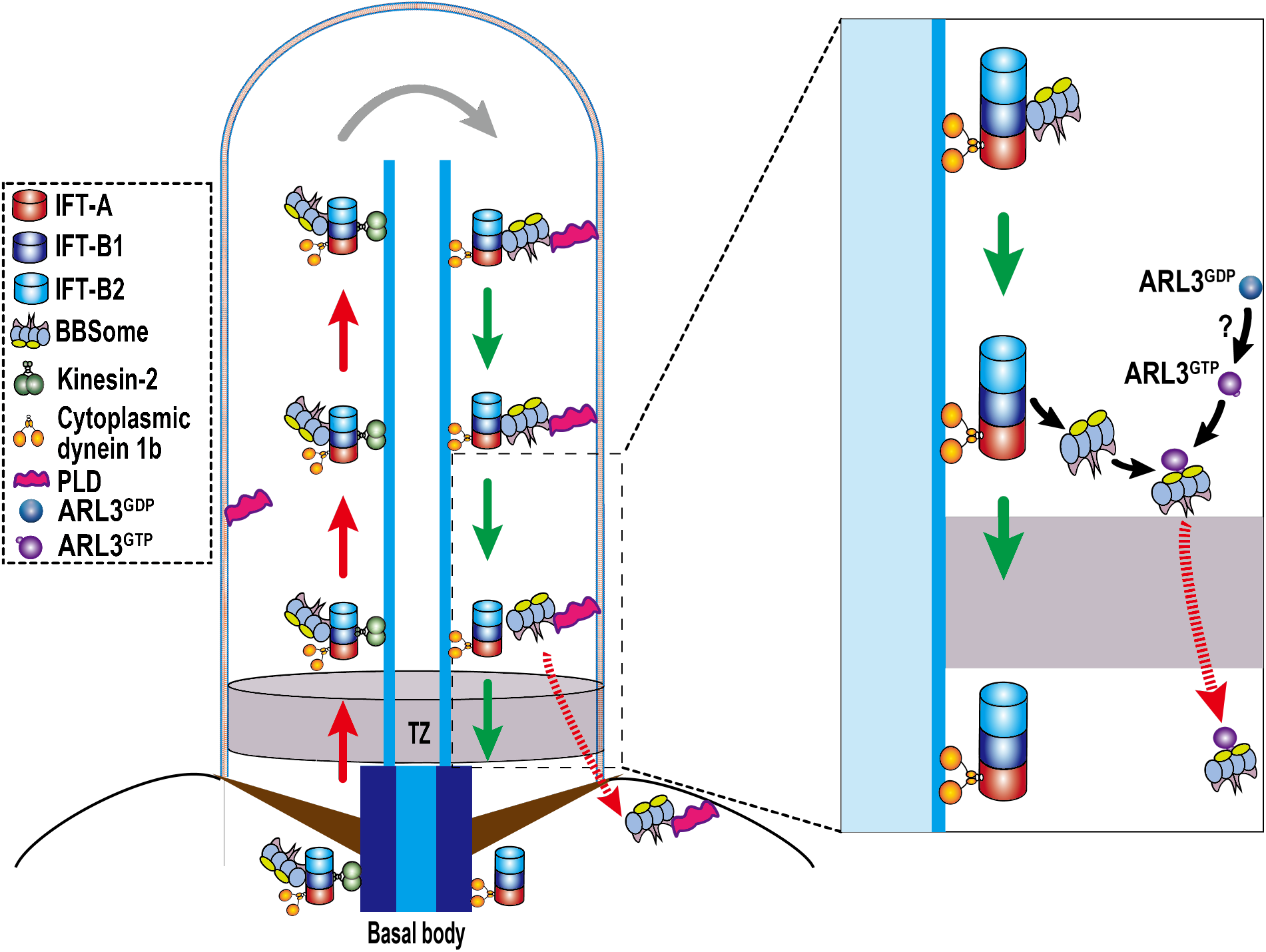
Hypothetical model for how ARL3 promotes outward PLD-laden BBSome diffusion through the TZ. ARL3^GDP^ diffuses into cilia and is activated to become ARL3^GTP^ by an unknown mechanism (?). Following the transportation from the ciliary tip to base, cargo (PLD) laden BBSomes shed from the retrograde IFT train at the proximal ciliary region right above the TZ and is bound to ARL3^GTP^ as a ARL3 effector. ARL3^GTP^ then recruits the cargo (PLD) laden BBSome to diffuse through the TZ for ciliary retrieval.

## Discussion

As a Arf-like small GTPase, *Chlamydomonas* ARL3 relies on GDP, its N-terminal amphipathic helix, and the glycine myristylation site at the second amino acid position to associate with the cell membrane, which is a prerequisite for ARL3 to diffuse into cilia, and to reside along the whole length of cilia by anchoring to the ciliary membrane. Following a rapid activation process in cilia, ARL3^GDP^ is converted to become ARL3^GTP^ for releasing from the ciliary membrane. Following ciliary cycling, the BBSome sheds from retrograde IFT at the proximal ciliary region right above the TZ. By acting as a major ARL3 effector, the cargo (PLD)-laden BBSome was then bound to and recruited by ARL3^GTP^ to diffuse through the TZ for ciliary retrieval (Fig. 7). Our data show that ARL3 maintains BBSome ciliary dynamics by moving it cross the diffusion barrier at the TZ and out of cilia, thus closing a gap in our understanding of how ARL3 affects cell behavior, e.g., phototaxis, of *C. reinhardtii*.

### How does ARL3 contribute to maintain BBSome dynamics in cilia**?**

The BBSome relies on IFT for maintaining its ciliary dynamics. During BBSome ciliary cycling, certain small GTPases contribute to maintain BBSome ciliary dynamics by mediating its coupling with IFT directly or indirectly. Our previous study identified RABL5/IFT22 binds ARL6/BBS3 to form an IFT22/BBS3 heterodimer in the cell body of *Chlamydomonas* cells and IFT22 binding is required for stabilizing BBS3 (Xue et al., 2020). As a major BBS3 effector, the BBSome is bound to and recruited by IFT22/BBS3 for targeting to the basal bodies (Jin et al., 2010; Xue et al., 2020). In such a way, IFT22/BBS3 controls BBSome amount available for entering cilia from the basal bodies, thus playing a critical role in maintaining BBSome ciliary dynamics (Xue et al., 2020). Upon reaching the ciliary tip via anterograde IFT, the BBSome disassembles first followed by reassembly, a process known as BBSome remodeling, before being able to load onto retrograde IFT trains for transporting to the ciliary base (Sun et al., 2021). During this process, RABL4/IFT27, by binding its stabilizing partner IFT25 to form an IFT25/27 heterodimer, cycles off IFT to promote BBSome reassembly (Liew et al., 2014; Sun et al., 2021; Wang et al., 2009). Therefore, IFT25/27 is critical for maintaining BBSome ciliary dynamics as it contributes to promote the BBSome to U turn at the ciliary tip. As for ARL3, it mimics ARL6/BBS3 to bind the membrane to diffuse into cilia and resides along the whole length of cilia by attaching to the ciliary membrane (Liu et al., 2021). Differing from ARL6/BBS3 that in a GTP-bound state binds the membrane, ARL3 relies on GDP for membrane binding (Liu et al., 2021). This is easy to understand as the cell may have developed an elaborated system to restrict ARL3 to bind its BBSome effector only at the proximal ciliary region right above the TZ where ARL3^GTP^ is supposed to concentrate (Fig. 2 F and 5 C). Upon reaching the proximal ciliary region right above the TZ via retrograde IFT, the BBSome drops off retrograde IFT via a mechanism that remains unknown yet, consistent with the mammalian BBSome behavior in cilia (Duan et al., 2021; Nachury, 2018; Ye et al., 2018). GTP loading then enables ARL3 to bind and recruit the IFT-detached BBSome to move cross the diffusion barrier at the TZ for ciliary retrieval. Therefore, ARL3 controls BBSome ciliary amount through mediating its diffusion through the TZ for ciliary removal, playing a critical role in maintaining BBSome ciliary dynamics.

### The BBSome is a ARL3 effector only when they both position to the proximal ciliary region

Cross ciliated species, PDE6D, UNC119A/B, and BART/BARTL1 are three types of ARL3 effectors known as carrier/solubilizing proteins for binding and shuttling cytoplasmic lipidated signaling protein cargoes into cilia (Linari et al., 1999; Lokaj et al., 2015; Wright et al., 2011). Once at the proximal ciliary region, ARL3 applies RP2 and ARL13b as its GAP and GEF, respectively, and is catalyzed to convert between being GTP- and GDP-bound (Gotthardt et al., 2015; Veltel et al., 2008; Zhang et al., 2016). This ARL3/effector/cargo cascade works efficiently for releasing the cytoplasmic lipidated cargoes to bind the ciliary membrane via their lipidated moieties. Reflecting its critical role in releasing signaling proteins downstream the transmembrane sensing receptors in cilia, ARL3 deficiency disrupts certain signal transduction pathway(s) to cause many diseases. i.e., inherited retinal degenerations (IRDs) (Fu et al., 2021; Ratnapriya et al., 2021). In this study, the BBSome was identified to be a major ARL3 effector in cilia, uncovering ARL3’s role in mediating ciliary signaling but via the BBSome pathway. Remarkably, ARL3 applies the BBSome as its effector only when they both position to the proximal ciliary region right above the TZ but not in the cell body, suggesting that ARL3 is intrinsically prevented from interacting with the BBSome for conducting GTPase/effector function in the cell body. This could be achieved simply by retaining ARL3 at a GDP-bound configuration in the cell body. Although the underlying mechanism of how cells restrain ARL3 in a GDP-bound state in the cell body compartment remains unknown thus far, ARL3 is indeed observed to exist in a GDP-bound state in the cell body, disabling it for BBSome binding in the cell body. It has been known that IFT22/BBS3 binds the BBSome via a direct interaction between the BBSome and BBS3 in the cell body and recruits the BBSome as a BBS3 effector to the basal bodies (Xue et al., 2020). This observation explained well why ARL3 even in a GTP-bound configuration fails to bind the BBSome in the cell body (Fig. 4 D and E). The cell may direct BBSome trafficking in different cellular compartments through applying distinct GTPase pathways.

### How is ARL3 activated to promote BBSome diffusion through the TZ?

ARL3 in a GDP-bound state enters cilia but relies on GTP for binding and recruiting the BBSome to diffuse through the TZ for ciliary retrieval, suggesting that ARL3 must have to convert from being GDP-bound to being GTP-bound in cilia. *Chlamydomonas* ARL13b has been identified as a ARL3 GEF to catalyze the conversion of ARL3^GDP^ to ARL3^GTP^ in cilia (Gotthardt et al., 2015). While ARL13b fits to the profile of the ARL3 GEF as it localizes to cilia and has in vitro GEF activity, functional correlation of ARL13b to ARL3 and structural studies for elucidating how ARL13b activates ARL3 at a molecular level were not recorded in the literature (ElMaghloob et al., 2021; Gotthardt et al., 2015). Most recently, ARL13b was found to be able to activate ARL3 but very weakly at physiological GTP:GDP levels and its stronger activation was achieved through applying BART as a so-called “co-GEF”, which stabilizes ARL3 to retain in a GTP-bound state (ElMaghloob et al., 2021). Unfortunately, knockout of human ARL13b alters BBSome ciliary dynamics but by reducing BBSome content in cilia, a result excluding ARL13b from promoting BBSome diffusion through the TZ for ciliary retrieval via ARL3 pathway (Fujisawa et al., 2021). Interestingly, although they both play critical roles in targeting INPP5E to the ciliary membrane in human cells, ARL3 and ARL13b instead participate in distinct steps of this event, again revealing their functional discrepancy in vivo (Fujisawa et al., 2021). These findings do not necessarily disapprove ARL13b for activating ARL3 as a possible functional ARL3 GEF in vivo, while ARL3/ARL13b cascade can be confidently excluded from promoting BBSome diffusion through the TZ for ciliary retrieval, at least in *C. reinhardtii*. This raised an interesting question, namely which factor other than ARL13b, if desirable, contributes to activate ARL3 specifically for promoting BBSome diffusion through the TZ. We currently had no answer for this question, while RABL2 deficiency causes BBSome to cease for moving out of cilia but to accumulate at the proximal region right above the TZ, the same BBSome intraciliary trafficking defect pattern as shown by ARL3 knock-out (Duan et al., 2021). This observation provides a clue, from the functional review, that RABL2 could likely be a ARL3 GEF in cilia, though his hypothesis remains to be confirmed.

### How does ARL3 mediate phototaxis?

*C. reinhardtii* cells apply the photoreceptors channelrhodopsin 1 (ChR1) and ChR2 to sense light for conducting phototaxis via ciliary beating (Berthold et al., 2008; Nagel et al., 2002; Nagel et al., 2003). Among these two sensory ion channels, ChR1 was identified as the major one, by residing in the eyespot, for sensing light to generate and transduce electrophysiological signal to cilia to direct their beating (Berthold et al., 2008). ChR1 was later shown to be able to target to cilia via an IFT-dependent manner, giving rise to a possibility that ChR1 might sense light for directing the cell to conduct phototaxis by residing in cilia but not the eyespot (Awasthi et al., 2016). We currently had no clue about if ChR1, like certain GPCRs (Ye et al., 2013; Ye et al., 2018), requires the BBSome for exporting out of cilia, while BBS mutants deprives the BBSome of being present in cilia, eventually disrupting phototaxis simply owing to biochemical defects of the ciliary membrane (Lechtreck et al., 2013; Lechtreck et al., 2009; Liu and Lechtreck, 2018). If the prediction that ChR1 ciliary positioning and its BBSome-dependent dynamics maintenance in cilia are prerequisites for *Chlamydomonas* cells to conduct phototaxis is correct, these defects may raise from abnormal ciliary buildup of ChR1 due to its disrupted export out of cilia in the absence of the BBSome. Therefore, it is most likely that ARL3 mediates phototaxis via controlling the BBSome-dependent ChR1 dynamics maintenance in cilia of *Chlamydomonas* cells, which deserves to be carefully investigated in the future.

## Materials and methods

### Antibodies, *Chlamydomonas* strains, and culture conditions

Rabbit-raised polyclonal antibodies including α-BBS1, α-BBS4, α-BBS5, α-BBS7, α-BBS8, α-IFT22, α-IFT38, α-IFT43, α-IFT46, α-IFT57, α-IFT70, α-IFT139, and α-PLD and mouse-raised α-IFT81 have been described previously and were listed in Table EV1 (Dong et al., 2017; Liu et al., 2021; Sun et al., 2021). Rabbit-originated antibodies against ARL3 and CEP290 were produced by Beijing Protein Innovation, LLC (Beijing). The monoclonal antibodies against YFP (mAbs 7.1 and 13.1, Roche), HA (mAb 3F10 clone, Roche), α-tubulin (mAb B512, Sigma), and acetylated-tubulin (mAb 6-11B-1, Sigma-Aldrich) were commercially bought (Table S1). *C. reinhardtii* strains including CC-125, CC-5325, and *arl3* (the CLiP mutant LMJ.RY0420.182282) were purchased from the *Chlamydomonas* Genetic Center at the University of Minnesota, Twin Cities, MN (http://www.chlamycollection.org). BBS8-null mutant *bbs8* has been reported previously (Sun and Pan, 2019). All the strains used in this study are listed in Table S2. If not otherwise specialized, strains were grown in Tris acetic acid phosphate (TAP) or minimal 1 (M1) medium in a continuous light with constant aeration at room temperature. Depending on a specific strain, cells were cultured with or without the addition of 20 µg/ml paromomycin (Sigma-Aldrich), 15 µg/ml bleomycin (Invitrogen), or both antibiotics with 10 µg/ml paromomycin and 5 µg/ml bleomycin.

### Vectors and cell line construction

ARL3 miRNA vector was created according to the method described previously (Hu et al., 2014). In brief, the miRNA sequences targeting the 3’-UTR region of *arl3* gene were designed using WMD3 software (http://wmd3.weigelworld.org) and were combined with the miRNA cre-MIR1157 (accession number MI0006219) to result in a 171-bp of ARL3 miRNA precursor sequence (Table S3). The ARL3 miRNA precursor sequence was synthesized by Genewiz (China) and ligated to the pHK263 plasmid (Hu et al., 2014), resulting in the ARL3 miRNA vector pMi-ARL3. Expression vectors were constructed on pBKS-gBBS3::HA::YFP-Ble that contained HA::YFP coding sequences followed immediately downstream by a sequence encoding the Rubisco 3’-UTR and the bleomycin cassette (*Ble*, zeocine resistant gene) (Dong et al., 2017). To generate ARL3::HA::YFP-expressing vector, a 3.4-kb ARL3 fragment consisting of the 1.0-kb promoter sequence and its coding region was amplified from genomic DNA by using primer pair (gARL3-FOR1 and gARL3-REV1) as listed in Table S3 and inserted into the *Xba*I and *Eco*RI sites of pBKS-gBBS3::HA::YFP-Ble, resulting in pBKS-gARL3::HA::YFP-Ble. To generate ARL3ΔN15::HA::YFP-expressing vector, two fragments were amplified by using primer pair (gARL3-FOR1 and gARL3ΔN15-REV; gARL3ΔN15-FOR and gARL3-REV1) as listed in Table S3 and inserted into the *Xba*I and *Eco*RI sites of pBKS-gBBS3::HA::YFP-Ble by three-way ligation, resulting in pBKS-gARL3ΔN15::HA::YFP-Ble. The desiring mutations including G2A, G2AT30N, G2AQ70L, T30N, and Q70L were introduced into pBKS-gARL3::HA::YFP-Ble and pBKS-gARL3ΔN15::HA::YFP-Ble by site-directed mutagenesis using the primer pairs (ARL3^G2A^-FOR and ARL3^G2A^-REV; ARL3^T30N^-FOR and ARL3^T30N^-REV; ARL3^Q70L^-FOR and ARL3^Q70L^-REV, respectively) as listed in Table S3. Afterward, the mutated DNAs were inserted into the *Xba*I and *Eco*RI sites of pBKS-gARL3::HA::YFP-Ble, resulting in pBKS-gARL3^T30N^::HA::YFP-Ble, pBKS-gARL3^Q70L^::HA::YFP-Ble, pBKS-gARL3^G2A^::HA::YFP-Ble, pBKS-gARL3^G2AT30N^::HA::YFP-Ble, pBKS-gARL3^G2AQ70L^::HA::YFP-Ble, pBKS-gARL3ΔN15^T30N^::HA::YFP-Ble, and pBKS-gARL3ΔN15^Q70L^::HA::YFP-Ble. To express IFT43::HA::YFP, a 2,800-bp DNA fragment composing of a 1,000-bp promoter and IFT43 coding sequence was amplified from genomic DNA by using the primer pair (gIFT43-FOR and gIFT43-REV) as listed in Table S3, and inserted into *Not*I and *Eco*RI sites of pBluescript II KS(+) vector, resulting in pBKS-gIFT43. Afterwards, gIFT43 sequence was cut from pBKS-gIFT43 by *Not*I and *Eco*RI sites, HA::YFP-Ble sequence was cut from pBKS-gARL3::HA::YFP-Ble by *Eco*RI and *Kpn*I, and inserted those two fragments into the *Not*I and *Kpn*I sites of pBluescript II KS(+) vector by three-way ligation, resulting in pBKS-gIFT43::HA::YFP-Ble. To express IFT38::YFP, a 3,875-bp DNA fragment composing of a 1,000-bp promoter sequence and the IFT38 coding sequence was amplified from genomic DNA using primer pair (gIFT38-FOR and gIFT38-REV) as listed in Table S3 and inserted into the *Bam*HI and *Eco*RI sites of pBluescript II KS(+) vector, resulting in pBKS-gIFT38. Next, IFT38 sequence was cut from pBKS-gIFT38, and inserted into the *Bam*HI and *Eco*RI sites of pBKS-gBBS5::YFP-Paro and pBKS-gBBS5::YFP-Ble, resulting in pBKS-gIFT38::YFP-Paro and pBKS-gIFT38::YFP-Ble, respectively. pBKS-gIFT22::HA::GFP-Paro and pBKS-gIFT22::HA::GFP-Ble have been described previously (Xue et al., 2020). To express IFT22::HA::YFP, the HA::GFP fragment was replaced with the HA::YFP fragment obtained from pBKS-gARL3::HA::YFP-Ble by *Eco*RI and *Xho*I digestion, resulting in pBKS-gIFT22::HA::YFP-Paro and pBKS-gIFT22::HA::YFP-Ble, respectively. To express BBS8::YFP, a 4,804-bp DNA fragment composing of 1,000-bp promoter and 3,804-bp BBS8 coding sequence was amplified from genomic DNA by using the primer pair (gBBS8-FOR and gBBS8-REV) as listed in Table S3, and inserted into *Xba*I and *Eco*RI sites of pBluescript II KS(+) vector , resulting in pBKS-gBBS8. Then, BBS8 sequence cut from pBKS-gBBS8 by *Xba*I and *Eco*RI sites and YFP-Ble sequence cut from pBKS-gBBS5::YFP-Ble by *Eco*RI and *Kpn*I sites were inserted into the *Xba*I and *Kpn*I sites of pBluescript II KS(+) vector by three-way ligation, resulting in pBKS-gBBS8::YFP-Ble. After the verification by direct nucleotide sequencing, the new constructs were transformed into *C. reinhardtii* strain by electroporation as described previously and screening of the positive transformants was done according to the method described previously (Xue et al., 2020). The screening of ARL3 miRNA cells was initiated by checking the cellular level of the target proteins through immunoblotting of whole cell extracts with ARL3 antibody. The miRNA strains showing a reduced level of the target proteins were selected for further phenotypic analysis.

For *arl3-bbs8* double mutant generation, 20 ml of each of *arl3-* and *bbs8*-null mutant cells were grown to a final concentration of 2×10^6^ cells/ml in M1 medium under continuous light at room temperature. The cells were collected by centrifugation at 1000 *×g* for 15 min and washed with M1-N (without nitrogen) medium. Afterwards, the cells were transferred to 20 ml of M1-N medium and aerated for 12 hrs under continuous light. After that, gametes of two opposite mating types were mixed in flask and incubated under light for 2 hrs. The cells sitting on the flask bottom were transferred to M1 plate containing 4% agar, air-dried, incubated overnight under continuous light, and then continued to be incubated for a week in the dark. The plates were then moved to incubate at -20°C for two days followed by culturing at room temperature under continuous light for at least ten days. The *arl3- and bbs8-*double null mutant was screened by detecting the loss of ARL3 and BBS8 proteins through immunoblotting of whole cell extracts with both ARL3 and BBS8 antibodies.

### Total RNA and genomic DNA manipulations

Genomic DNA of *Chlamydomonas* cells was extracted and purified using a Wizard® Genomic DNA Purification Kit (Promega, Beijing) following the kit’s protocol. To characterize the *arl3* cell (the CLiP mutant LMJ.RY0420.182282) at the genomic level, 20 ng of genomic DNA was applied as PCR template to amplify ARL3 genomic sequence. The PCR reactions were performed at 95°C for 5 min followed by 30 cycles of 95 °C for 20 sec, 61°C for 20 sec, and 72°C for 5 min with the primer pair gARL3-FOR1 and gARL3-REV1 as listed in Table S3. Total RNA of *Chlamydomonas* cells was extracted and purified according to our protocol reported previously (Dong et al., 2017). Five micrograms of RNA were reverse transcribed at 42 °C for 1 h using M-MLV Reverse Transcriptase (Promega) and oligo(T)18 primers (Takara). ARL3 cDNA and cDNAs encoding the six BBS proteins BBS1, BBS2, BBS4, BBS5, BBS7, and BBS8 were amplified by PCR using primer pairs as listed in Table S3. The PCR reactions were performed at 95 °C for 5 min followed by 30 cycles of 95°C for 20 sec, 61°C for 20 sec, and 72°C for 4 min.

### Ciliary length measurement

*Chlamydomonas* cells growing to a concentration of ∼10^7^ cells were collected and placed onto the surface of glass slides and covered by cover glass. The cells were observed under an IX83 inverted fluorescent microscopy at 100× amplification. Phase contrast images were then taken for ciliary length analysis. Ciliary length was measured by using ImageJ (version 1.42g, National Institutes of Health) according to our protocol reported previously (Fan et al., 2010). The data was processed with GraphPad Prism 8.30 (GraphPad Software). For each strain, a total of 20 cilia were measured.

### Isolation of cilia and cell bodies

Isolation of cilia and cell bodies was performed according to our protocol reported previously (Fan et al., 2010). In brief, *Chlamydomonas* cells were grown in 10 liters of TAP medium to a final density of 10^8^ cells/ml, collected by centrifugation at 1000 *×g* for 15 min, and suspended in 150 ml of TAP (pH7.4). Cells were incubated for 2 h under strong light with bubbling before 0.5 M acetic acid was added to adjust the pH value to 4.5 for cell deciliation. Afterwards, 0.5 M KOH was added to adjust the pH value to 7.4. Cell body pellets and cilia in the supernatant were collected separately after centrifugation at 600 *×g* at 4°C for 5 min. To avoid the possible cell body contamination, cilia were repeatedly washed with HMDEKN buffer (30 mM Hepes [pH 7.4], 5 mM MgSO_4_, 1 mM DTT, 0.5 mM EGTA, 25 mM KCl, 125 mM NaCl) by centrifugation at 12,000 ×*g* for 10 min until the green color disappeared completely. All the experiments were done at 4°C.

### Preparation of ciliary fractions

Ciliary fractions were prepared according to our protocol reported previously (Liu et al., 2021). In brief, cell body-depleted cilia were dissolved in HMDEKN buffer supplemented with protein inhibitors (PI) (1 mM PMSF, 50 μg/ml soy-bean trypsin inhibitor, 1 μg/ml pepstatin A, 2 μg/ml aprotinin, and 1 μg/ml leupeptin) and frozen in liquid nitrogen. After three cycles of frozen-and-thaw, the solution was centrifugated at 12,000 ×g at 4°C for 15 min and the ciliary matrix fraction was collected as the supernatant. The pellets were then dissolved in HMEDKN (see above) buffer containing 0.5% nonidet P-40 (NP-40) and stayed on ice for 15 min. After centrifugation at 12,000 ×g at 4°C for 10 min, the supernatant and pellet were collected as membrane and axonemal fractions, respectively.

### Sucrose density gradient centrifugation assay

Ciliary samples were analyzed by sucrose density gradient centrifugation according to our protocol reported previously (Sun et al., 2021). Briefly, linear 12 ml of 10-25% sucrose density gradients were prepared in HMDEKN buffer supplemented with PI (see above) and 1% NP-40. Ciliary extracts were frozen and thawed for three cycles by using liquid nitrogen and centrifuged at 12,000 *×g* at 4 for 10 min. After the non-soluble debris was removed, 700 μl of samples were loaded on the top of the gradients and separated at 38,000 *×g* for 14 hrs in a SW41Ti rotor (Beckman Coulter). After the gradients were fractioned into 24 0.5 ml aliquots, 20 μl of each fraction was loaded into SDS-PAGE gel for electrophoresis and analyzed by immunoblotting as described below. The centrifugation was done at 4°C.

### Immunoblotting

Whole cell, cell body, and ciliary samples were prepared for immunoblotting according to our protocol reported previously (Dong et al., 2017). If not otherwise specified, 20 µg of total protein from each sample was loaded in the 12% SDS-PAGE gel for electrophoresis. Primary and secondary antibodies were diluted for immunoblotting with a ratio as shown in Table S1. ImageJ software (version 1.42g, National Institutes of Health) was applied for quantifying the target proteins by measuring the immunoblot intensity according to our protocol reported previously (Xue et al., 2020). The immunoblot intensity was normalized to the intensity of a loading control protein.

### Immunoprecipitation

Immunoprecipitation was performed according to our protocol reported previously (Liu et al., 2021). In brief, cell body and ciliary samples isolated from cells expressing HA::YFP, HA::YFP-tagged ARL3, or its variants were resuspended in HMDEKN or DTT-deprived HMEKN buffer supplemented with protein inhibitors (see above). The samples were lysed by adding NP-40 to a final concentration of 1% followed by centrifugation at 14,000 ×*g*, 4°C for 10 min. Afterward, the supernatants were collected for agitating with 5% BSA-pretreated camel anti-YFP antibody-conjugated agarose beads (V-nanoab Biotechnology) for 2 hrs at 4°C. After continuous washing with HMEKN buffer, HMEK buffer containing 50 mM NaCl, and HMEK without containing NaCl, the beads were collected by centrifugation at 2,500 ×*g* for 2 min, mixed with Laemmli SDS sample buffer, and boiled for 5 min before centrifugation at 2,500 *×g* for 5 min. The immunoprecipitants in the supernatants were collected for immunoblotting analysis as described above.

### Protein-liposome binding assay

Liposomes were prepared according to the protocol reported previously (Jin et al., 2010). The ARL3 cDNA was inserted into the *EcoR*I and *Xho*I sites of pET-28a (Novagen) to result in pET-28a-cARL3. The desiring deletion of N-terminal 15 amino acid residues and the mutations G2A, Q70L, and T30N were introduced into ARL3 by regular PCR or site-directed mutagenesis using the primer pairs as listed in Table S3, resulting plasmids pET-28a-cARL3^Q70L^, pET-28a-cARL3^T30N^, pET-28a-cARL3ΔN15, pET-28a-cARL3ΔN15^Q70L^, pET-28a-cARL3ΔN15^T30N^, pET-28a-cARL3^G2A^, pET-28a-cARL3^G2AQ70L^, and pET-28a-cARL3^G2AT30N^, respectively. After these plasmids were transformed into the *Escherichia coli* strain BL21(DE3), the bacterially expressed 6×His tagged ARL3 and its variants were purified with Ni Sepharose^TM^ 6 Fast Flow beads (GE Healthcare) and cleaved with thrombin (Solarbio) for 1 hr at 37°C to get rid of the 6×His tag according to our previous report (Xue et al., 2020). ARL3 association with liposome was conducted in binding buffer (50 mM Tris, pH 7.5, 150 mM NaCl, 0.05% NP-40). In brief, 1 µg of the purified ARL3 (in the presence of 100 mM of GTPγS or of GDP) or its variants protein, 20 µl of 1 mM PolyPIPosomes^TM^ (Echelon Biosciences), and 1 ml of binding buffer were mixed, rotated for 10 min at room temperature, and then centrifuged at 12,000 ×g for 10 min. The liposome pellet was washed in 1 ml of binding buffer by centrifugation for three times. Afterward, equal portions of the resulting pellets were mixed with Laemmli SDS sample buffer and boiled for 5 min before resolving by SDS-PAGE and immunoblotting with α-ARL3.

### Fixed imaging

Immunofluorescence staining was performed according to our protocol reported previously (Wang et al., 2009). The primary antibodies against CEP290, BBS8, IFT46, IFT81, YFP, HA, and the secondary antibodies including Alexa-Fluor594-conjugated goat anti-rabbit, Alexa-Fluor488-conjugated goat anti-mouse, and Alexa-Fluor488-conjugated goat anti-rat (Molecular Probes, Eugene, OR) were listed in Table S1 with their suggested dilutions. Images were captured with an IX83 inverted fluorescent microscopy (Olympus) equipped with a back illuminated scientific CMOS camera (Prime 95B, Photometrics), a 100×/1.40 NA oil objective lens (Olympus), and 488-nm and 561-nm lasers from Coherent OBIS Laser Module. All images were acquired and processed with CellSens Dimension (version 2.1, Olympus).

### Live-cell imaging

Total internal reflection fluorescence (TIRF) microscopy was applied to visualize the motility of YFP-tagged IFT43, IFT22, IFT38, IFT46, BBS8, and ARL3 and its variants in cilia. YFP-tagged proteins was imaged at ∼15 frames per second (fps) using an IX83 inverted fluorescent microscopy (Olympus) equipped with a through-the-objective TIRF system, a 100×/1.49 NA TIRF oil immersion objective len (Olympus), a back illuminated scientific CMOS camera (Prime 95B, Photometrics), and 488-nm laser from Coherent OBIS Laser Module as detailed previously (Xue et al., 2020).

### Kymogram analysis

Kymography was generated according to our protocol reported previously (Dong et al., 2017). In brief, YFP-tagged IFT, BBSome and ARL3 proteins were imaged with TIRF (15 Hz) for ∼20 s. The videos obtained were processed with CellSens Dimension (version 2.1, Olympus) for generating kymographs. In kymographs, lines drawn along the long axis of the cilium (“the leg”) and the processive movement (“the hypotenuse”) represent the anterograde and retrograde IFT tracks. The angle of the lines was measured (“the included angle”). Comparison of the leg and the hypotenuse was applied for quantifying the frequency of an IFT- or BBS-containing train. Comparison of the hypotenuse and the included angle was applied for measuring the velocity of IFT and BBSome trains. KymographClear was applied to deconvolve retrograde from anterograde trains for clearly showing BBSome diffusion through the transition zone (e.g., Fig. 5E) (Mangeol et al., 2016).

### Protein-protein interaction assay

The cDNA encoding BBS9 was synthesized by Genewiz (China). The cDNAs encoding BBS1, BBS2, BBS4, BBS5, BBS7, BBS8, and BBS9 were inserted into the *Bam*HI and *Hin*d III sites of pMal-C2x (Nova Lifetech) to generate pMal-C2x-cBBS1, pMal-C2x-cBBS2, pMal-C2x-cBBS4, pMal-C2x-cBBS5, pMal-C2x-cBBS7, pMal-C2x-cBBS8, and pMal-C2x-cBBS9. After these plasmids and the empty pMal-C2x plasmid were transformed into the *E. coli* strain BL21(DE3), the bacterially expressed N-terminal MBP-tagged BBS proteins were purified with Dextrin Sepharose^TM^ High Performance with MBP-tagged protein purification resin (GE Healthcare). One hundred micrograms of MBP and the MBP-tagged BBS proteins were individually mixed with 100 µg of bacterially expressed ARL3^Q70L^ or ARL3^T30N^ (see above) to form a combination of 16 reactions. After incubated for 2 hrs at room temperature, the mixtures were purified with Dextrin Sepharose^TM^ High Performance MBP-tagged protein purification resin (GE Healthcare). Ten micrograms of proteins from elutes was resolved on 12% SDS-PAGE gels and visualized with Coomassie blue staining. Immunoblotting assay was also performed to verify the interaction between BBS proteins and ARL3 variants with α-ARL3.

### Population phototaxis assay

Population phototaxis assays were performed on *Chlamydomonas* cells according to the protocol reported previously (Liu and Lechtreck, 2018). In brief, *Chlamydomonas* cells growing to a concentration of ∼10^7^ cells were harvested and 100 μl of the cell suspension were placed into the surface of Petri dishes of 3.5-cm diameter (706001; Wuxi NEST Biotech.) containing solid TAP medium. Afterwards, the cells were illuminated with a flashlight from one side for 4 min. Images were continuously taken once every two minutes with a standard digital camera (Nikon A70).

### Single-cell motion assay

Single-cell motion assay was performed on *Chlamydomonas* cells according to the protocol reported previously (Liu and Lechtreck, 2018). Briefly, 20 μl of cell suspensions as obtained above were placed on superfrost™ plus microscope slides (12-550-15; Fisher Brand) and observed using an inverted microscope (IX83, Olympus) under non-phototactic red light illumination. The cells were then illuminated for 5s with phototactic active green light. Afterwards, five images were continuously taken once every 0.3s using a back illuminated scientific CMOS camera (Prime 95B, Photometrics). The five sequential images each displayed in a different color were merged in ImageJ (version 1.42g, National Institutes of Health) to show the swimming tracks of single cells, allowing us to determine the angle and the direction of a cell’s movements. Excel for Mac (version 16.52) were applied to analyze the data and generate polar histograms with 60° bins.

### Statistical analysis

Statistical analysis was done with GraphPad Prism 8.30 (GraphPad Software). For comparisons on velocities and frequencies of the YFP- and HA::YFP-labeled proteins and ciliary length measurement, one-sample unpaired student *t*-test was used on samples. The data were presented as mean ± S.D. n.s. represents non-significance.

### Data availability

Data supporting the findings of this study were contained within this paper and the supplementary files.

## Acknowledgments

Research reported in this publication was supported by National Natural Science Foundation of China (32070698 to Z-C.F. and 32100541 to B.X) and China Postdoctoral Science Foundation (2021M702457 to Y-X.L. and 2021M692403 to B.X). The founders have no role in study design, data collection and analysis, decision to publish, or preparation of the manuscript.

## Author contributions

Z-C.F. conceived and directed the project. Y-X.L. performed major experiments. Y-X.L. and W-Y.S. performed immunostaining assays. B.X. screened the *bbs8*::BBS8::YFP cell. R-K.Z. produced the ARL3 antibody. W-J.L. assisted in isolating and purifying cilia. Y-X.L., X.X. and Z-C.F. analyzed the data. Z-C.F. interpreted and wrote the paper.

## References

Abd-El-Barr, M.M., K. Sykoudis, S. Andrabi, E.R. Eichers, M.E. Pennesi, P.L. Tan, J.H. Wilson, N. Katsanis, J.R. Lupski, and S.M. Wu. 2007. Impaired photoreceptor protein transport and synaptic transmission in a mouse model of Bardet-Biedl syndrome. Vision Res. 47:3394–3407.

Amor, J.C., D.H. Harrison, R.A. Kahn, and D. Ringe. 1994. Structure of the human ADP-ribosylation factor 1 complexed with GDP. Nature. 372:704–708.

Avidor-Reiss, T., A.M. Maer, E. Koundakjian, A. Polyanovsky, T. Keil, S. Subramaniam, and C.S. Zuker. 2004. Decoding cilia function: defining specialized genes required for compartmentalized cilia biogenesis. Cell. 117:527–539.

Awasthi, M., P. Ranjan, K. Sharma, S.K. Veetil, and S. Kateriya. 2016. The trafficking of bacterial type rhodopsins into the Chlamydomonas eyespot and flagella is IFT mediated. Sci Rep. 6:34646.

Berthold, P., S.P. Tsunoda, O.P. Ernst, W. Mages, D. Gradmann, and P. Hegemann. 2008. Channelrhodopsin-1 initiates phototaxis and photophobic responses in chlamydomonas by immediate light-induced depolarization. The Plant cell. 20:1665–1677.

Cuvillier, A., F. Redon, J.C. Antoine, P. Chardin, T. DeVos, and G. Merlin. 2000. LdARL-3A, a Leishmania promastigote-specific ADP-ribosylation factor-like protein, is essential for flagellum integrity. J Cell Sci. 113 ( Pt 11):2065–2074.

Dateyama, I., Y. Sugihara, S. Chiba, R. Ota, R. Nakagawa, T. Kobayashi, and H. Itoh. 2019. RABL2 positively controls localization of GPCRs in mammalian primary cilia. J Cell Sci. 132.

Dong, B., S. Wu, J. Wang, Y.X. Liu, Z. Peng, D.M. Meng, K. Huang, M. Wu, and Z.C. Fan. 2017. Chlamydomonas IFT25 is dispensable for flagellar assembly but required to export the BBSome from flagella. Biol Open. 6:1680–1691.

Duan, S., H. Li, Y. Zhang, S. Yang, Y. Chen, B. Qiu, C. Huang, J. Wang, J. Li, X. Zhu, and X. Yan. 2021. Rabl2 GTP hydrolysis licenses BBSome-mediated export to fine-tune ciliary signaling. EMBO J. 40:e105499.

Efimenko, E., K. Bubb, H.Y. Mak, T. Holzman, M.R. Leroux, G. Ruvkun, J.H. Thomas, and P. Swoboda. 2005. Analysis of xbx genes in C. elegans. Development. 132:1923–1934.

ElMaghloob, Y., B. Sot, M.J. McIlwraith, E. Garcia, T. Yelland, and S. Ismail. 2021. ARL3 activation requires the co-GEF BART and effector-mediated turnover. Elife. 10:e64624.

Fan, Z.-C., R.H. Behal, S. Geimer, Z. Wang, S.M. Williamson, H. Zhang, D.G. Cole, and H. Qin. 2010. Chlamydomonas IFT70/CrDYF-1 Is a Core Component of IFT Particle Complex B and Is Required for Flagellar Assembly. Mol Biol Cell. 21:2696–2706.

Fansa, E.K., S.K. Kosling, E. Zent, A. Wittinghofer, and S. Ismail. 2016. PDE6delta-mediated sorting of INPP5E into the cilium is determined by cargo-carrier affinity. Nat Commun. 7:11366.

Fu, L., Y. Li, S. Yao, Q. Guo, Y. You, X. Zhu, and B. Lei. 2021. Autosomal Recessive Rod-Cone Dystrophy Associated With Compound Heterozygous Variants in ARL3 Gene. Frontiers in Cell and Developmental Biology. 9.

Fujisawa, S., H. Qiu, S. Nozaki, S. Chiba, Y. Katoh, and K. Nakayama. 2021. ARL3 and ARL13B GTPases participate in distinct steps of INPP5E targeting to the ciliary membrane. Biol Open.

Gillingham, A.K., and S. Munro. 2007. The Small G Proteins of the Arf Family and Their Regulators. Annual Review of Cell and Developmental Biology. 23:579–611.

Goetz, S.C., and K.V. Anderson. 2010. The primary cilium: a signalling centre during vertebrate development. Nat Rev Genet. 11:331–344.

Gotthardt, K., M. Lokaj, C. Koerner, N. Falk, A. Giessl, and A. Wittinghofer. 2015. A G-protein activation cascade from Arl13B to Arl3 and implications for ciliary targeting of lipidated proteins. Elife. 4.

Hanke-Gogokhia, C., Z. Wu, C.D. Gerstner, J.M. Frederick, H. Zhang, and W. Baehr. 2016. Arf-like Protein 3 (ARL3) Regulates Protein Trafficking and Ciliogenesis in Mouse Photoreceptors. Journal of Biological Chemistry. 291:7142–7155.

Hildebrandt, F., T. Benzing, and N. Katsanis. 2011. Ciliopathies. N Engl J Med. 364:1533–1543.

Hu, J., X. Deng, N. Shao, G. Wang, and K. Huang. 2014. Rapid construction and screening of artificial microRNA systems in Chlamydomonas reinhardtii. Plant J. 79:1052–1064.

Ismail, S.A., Y.X. Chen, M. Miertzschke, I.R. Vetter, C. Koerner, and A. Wittinghofer. 2012. Structural basis for Arl3-specific release of myristoylated ciliary cargo from UNC119. EMBO J. 31:4085–4094.

Jin, H., S.R. White, T. Shida, S. Schulz, M. Aguiar, S.P. Gygi, J.F. Bazan, and M.V. Nachury. 2010. The conserved Bardet-Biedl syndrome proteins assemble a coat that traffics membrane proteins to cilia. Cell. 141:1208–1219.

Lechtreck, K.F., J.M. Brown, J.L. Sampaio, J.M. Craft, A. Shevchenko, J.E. Evans, and G.B. Witman. 2013. Cycling of the signaling protein phospholipase D through cilia requires the BBSome only for the export phase. J Cell Biol. 201:249–261.

Lechtreck, K.F., E.C. Johnson, T. Sakai, D. Cochran, B.A. Ballif, J. Rush, G.J. Pazour, M. Ikebe, and G.B. Witman. 2009. The Chlamydomonas reinhardtii BBSome is an IFT cargo required for export of specific signaling proteins from flagella. J Cell Biol. 187:1117–1132.

Li, N., and W. Baehr. 1998. Expression and characterization of human PDEdelta and its Caenorhabditis elegans ortholog CEdelta. FEBS Lett. 440:454–457.

Li, Y., Q. Wei, Y. Zhang, K. Ling, and J. Hu. 2010. The small GTPases ARL-13 and ARL-3 coordinate intraflagellar transport and ciliogenesis. J Cell Biol. 189:1039–1051.

Liew, G.M., F. Ye, A.R. Nager, J.P. Murphy, J.S. Lee, M. Aguiar, D.K. Breslow, S.P. Gygi, and M.V. Nachury. 2014. The intraflagellar transport protein IFT27 promotes BBSome exit from cilia through the GTPase ARL6/BBS3. Dev Cell. 31:265–278.

Linari, M., M. Hanzal-Bayer, and J. Becker. 1999. The delta subunit of rod specific cyclic GMP phosphodiesterase, PDE delta, interacts with the Arf-like protein Arl3 in a GTP specific manner. FEBS Lett. 458:55–59.

Liu, P., and K.F. Lechtreck. 2018. The Bardet–Biedl syndrome protein complex is an adapter expanding the cargo range of intraflagellar transport trains for ciliary export. Proc Natl Acad Sci U S A. 115:E934–E943.

Liu, P., X. Lou, J.L. Wingfield, J. Lin, D. Nicastro, and K. Lechtreck. 2020. Chlamydomonas PKD2 organizes mastigonemes, hair-like glycoprotein polymers on cilia. J Cell Biol. 219.

Liu, Y., R.A. Kahn, and J.H. Prestegard. 2010. Dynamic structure of membrane-anchored Arf*GTP. Nat Struct Mol Biol. 17:876–881.

Liu, Y.-X., B. Xue, W.-Y. Sun, J.L. Wingfield, J. Sun, M. Wu, K.F. Lechtreck, Z. Wu, and Z.-C. Fan. 2021. Bardet-Biedl syndrome 3 protein promotes ciliary exit of the signaling protein phospholipase D via the BBSome. Elife. 10:e59119.

Lokaj, M., S.K. Kosling, C. Koerner, S.M. Lange, S.E. van Beersum, J. van Reeuwijk, R. Roepman, N. Horn, M. Ueffing, K. Boldt, and A. Wittinghofer. 2015. The Interaction of CCDC104/BARTL1 with Arl3 and Implications for Ciliary Function. Structure. 23:2122–2132.

Loktev, A.V., Q. Zhang, J.S. Beck, C.C. Searby, T.E. Scheetz, J.F. Bazan, D.C. Slusarski, V.C. Sheffield, P.K. Jackson, and M.V. Nachury. 2008. A BBSome subunit links ciliogenesis, microtubule stability, and acetylation. Dev Cell. 15:854–865.

Lv, B., L. Wan, M. Taschner, X. Cheng, E. Lorentzen, and K. Huang. 2017. Intraflagellar transport protein IFT52 recruits IFT46 to the basal body and flagella. J Cell Sci. 130:1662–1674.

Mangeol, P., B. Prevo, and E.J. Peterman. 2016. KymographClear and KymographDirect: two tools for the automated quantitative analysis of molecular and cellular dynamics using kymographs. Mol Biol Cell. 27:1948–1957.

Mourão, A., A.R. Nager, M.V. Nachury, and E. Lorentzen. 2014. Structural basis for membrane targeting of the BBSome by ARL6. Nat Struct Mol Biol. 21:1035.

Nachury, M.V. 2018. The molecular machines that traffic signaling receptors into and out of cilia. Curr Opin Cell Biol. 51:124–131.

Nachury, M.V., A.V. Loktev, Q. Zhang, C.J. Westlake, J. Peranen, A. Merdes, D.C. Slusarski, R.H. Scheller, J.F. Bazan, V.C. Sheffield, and P.K. Jackson. 2007. A core complex of BBS proteins cooperates with the GTPase Rab8 to promote ciliary membrane biogenesis. Cell. 129:1201–1213.

Nachury, M.V., and D.U. Mick. 2019. Establishing and regulating the composition of cilia for signal transduction. Nat Rev Mol Cell Biol. 20:389–405.

Nagel, G., D. Ollig, M. Fuhrmann, S. Kateriya, A.M. Musti, E. Bamberg, and P. Hegemann. 2002. Channelrhodopsin-1: a light-gated proton channel in green algae. Science. 296:2395–2398.

Nagel, G., T. Szellas, W. Huhn, S. Kateriya, N. Adeishvili, P. Berthold, D. Ollig, P. Hegemann, and E. Bamberg. 2003. Channelrhodopsin-2, a directly light-gated cation-selective membrane channel. Proceedings of the National Academy of Sciences of the United States of America. 100:13940–13945.

Nakayama, K., and Y. Katoh. 2020. Architecture of the IFT ciliary trafficking machinery and interplay between its components. Critical Reviews in Biochemistry and Molecular Biology:1–18.

Nishimura, D.Y., M. Fath, R.F. Mullins, C. Searby, M. Andrews, R. Davis, J.L. Andorf, K. Mykytyn, R.E. Swiderski, B. Yang, R. Carmi, E.M. Stone, and V.C. Sheffield. 2004. Bbs2-null mice have neurosensory deficits, a defect in social dominance, and retinopathy associated with mislocalization of rhodopsin. Proc Natl Acad Sci U S A. 101:16588–16593.

Nozaki, S., R.F. Castro Araya, Y. Katoh, and K. Nakayama. 2019. Requirement of IFT-B-BBSome complex interaction in export of GPR161 from cilia. Biol Open.

Pazour, G.J., N. Agrin, J. Leszyk, and G.B. Witman. 2005. Proteomic analysis of a eukaryotic cilium. J Cell Biol. 170:103–113.

Ratnapriya, R., S.G. Jacobson, A.V. Cideciyan, M.A. English, A.J. Roman, A. Sumaroka, R. Sheplock, and A. Swaroop. 2021. A Novel ARL3 Gene Mutation Associated With Autosomal Dominant Retinal Degeneration. Frontiers in Cell and Developmental Biology. 9.

Sahin, A., B. Espiau, E. Tetaud, A. Cuvillier, L. Lartigue, A. Ambit, D. Robinson, and G. Merlin. 2008. The Leishmania ARL-1 and Golgi traffic. PloS one. 3:e1620.

Schneider, L., C.A. Clement, S.C. Teilmann, G.J. Pazour, E.K. Hoffmann, P. Satir, and S.T. Christensen. 2005. PDGFRalphaalpha signaling is regulated through the primary cilium in fibroblasts. Curr Biol. 15:1861–1866.

Schrick, J.J., P. Vogel, A. Abuin, B. Hampton, and D.S. Rice. 2006. ADP-ribosylation factor-like 3 is involved in kidney and photoreceptor development. Am J Pathol. 168:1288–1298.

Singla, V., and J.F. Reiter. 2006. The primary cilium as the cell’s antenna: signaling at a sensory organelle. Science. 313:629–633.

Su, X., K. Driscoll, G. Yao, A. Raed, M. Wu, P.L. Beales, and J. Zhou. 2014. Bardet–Biedl syndrome proteins 1 and 3 regulate the ciliary trafficking of polycystic kidney disease 1 protein. Hum Mol Genet. 23:5441–5451.

Sun, L., and J. Pan. 2019. Bardet-Biedl syndrome protein-8 is involved in flagellar membrane protein transport in Chlamydomonas reinhardtii. Sheng Wu Gong Cheng Xue Bao. 35:133–141.

Sun, W.-Y., B. Xue, Y.-X. Liu, R.-K. Zhang, R.-C. Li, W. Xin, M. Wu, and Z.-C. Fan. 2021.Chlamydomonas LZTFL1 mediates phototaxis via controlling BBSome recruitment to the basal body and its reassembly at the ciliary tip. Proceedings of the National Academy of Sciences. 118:e2101590118.

Taschner, M., and E. Lorentzen. 2016. The Intraflagellar Transport Machinery. Cold Spring Harb Perspect Biol. 8.

Thomas, S., K.J. Wright, S. Le Corre, A. Micalizzi, M. Romani, A. Abhyankar, J. Saada, I. Perrault, J. Amiel, J. Litzler, E. Filhol, N. Elkhartoufi, M. Kwong, J.L. Casanova, N. Boddaert, W. Baehr, S. Lyonnet, A. Munnich, L. Burglen, N. Chassaing, F. Encha-Ravazi, M. Vekemans, J.G. Gleeson, E.M. Valente, P.K. Jackson, I.A. Drummond, S. Saunier, and T. Attie-Bitach. 2014. A homozygous PDE6D mutation in Joubert syndrome impairs targeting of farnesylated INPP5E protein to the primary cilium. Hum Mutat. 35:137–146.

Vaughan, M., and J. Moss. 1997. Activation of toxin ADP-ribosyltransferases by the family of ADP-ribosylation factors. Adv Exp Med Biol. 419:315–320.

Veltel, S., A. Kravchenko, S. Ismail, and A. Wittinghofer. 2008. Specificity of Arl2/Arl3 signaling is mediated by a ternary Arl3-effector-GAP complex. FEBS Lett. 582:2501–2507.

Wang, Z., Z.-C. Fan, S.M. Williamson, and H. Qin. 2009. Intraflagellar transport (IFT) protein IFT25 is a phosphoprotein component of IFT complex B and physically interacts with IFT27 in Chlamydomonas. PLoS One. 4:e5384.

Watzlich, D., I. Vetter, K. Gotthardt, M. Miertzschke, Y.X. Chen, A. Wittinghofer, and S. Ismail. 2013. The interplay between RPGR, PDEdelta and Arl2/3 regulate the ciliary targeting of farnesylated cargo. EMBO Rep. 14:465–472.

Wright, K.J., L.M. Baye, A. Olivier-Mason, S. Mukhopadhyay, L. Sang, M. Kwong, W. Wang, P.R. Pretorius, V.C. Sheffield, P. Sengupta, D.C. Slusarski, and P.K. Jackson. 2011. An ARL3– UNC119–RP2 GTPase cycle targets myristoylated NPHP3 to the primary cilium. Genes & Development. 25:2347–2360.

Xue, B., Y.-X. Liu, B. Dong, J.L. Wingfield, M. Wu, J. Sun, K.F. Lechtreck, and Z.-C. Fan. 2020. Intraflagellar transport protein RABL5/IFT22 recruits the BBSome to the basal body through the GTPase ARL6/BBS3. Proc Natl Acad Sci U S A. 117:2496–2505.

Ye, F., D.K. Breslow, E.F. Koslover, A.J. Spakowitz, W.J. Nelson, and M.V. Nachury. 2013. Single molecule imaging reveals a major role for diffusion in the exploration of ciliary space by signaling receptors. Elife. 2:e00654.

Ye, F., A.R. Nager, and M.V. Nachury. 2018. BBSome trains remove activated GPCRs from cilia by enabling passage through the transition zone. J Cell Biol. 217:1847–1868.

Yeh, C., A. Li, J.Z. Chuang, M. Saito, A. Caceres, and C.H. Sung. 2013. IGF-1 activates a cilium-localized noncanonical Gbetagamma signaling pathway that regulates cell-cycle progression. Dev Cell. 26:358–368.

Zhang, H., R. Constantine, S. Vorobiev, Y. Chen, J. Seetharaman, Y.J. Huang, R. Xiao, G.T. Montelione, C.D. Gerstner, M.W. Davis, G. Inana, F.G. Whitby, E.M. Jorgensen, C.P. Hill, L. Tong, and W. Baehr. 2011a. UNC119 is required for G protein trafficking in sensory neurons. Nat Neurosci. 14:874–880.

Zhang, H., S. Li, T. Doan, F. Rieke, P.B. Detwiler, J.M. Frederick, and W. Baehr. 2007. Deletion of PrBP/delta impedes transport of GRK1 and PDE6 catalytic subunits to photoreceptor outer segments. Proc Natl Acad Sci U S A. 104:8857–8862.

Zhang, H., X.H. Liu, K. Zhang, C.K. Chen, J.M. Frederick, G.D. Prestwich, and W. Baehr. 2004. Photoreceptor cGMP phosphodiesterase delta subunit (PDEdelta) functions as a prenyl-binding protein. J Biol Chem. 279:407–413.

Zhang, Q., J. Hu, and K. Ling. 2013. Molecular views of Arf-like small GTPases in cilia and ciliopathies. Exp Cell Res. 319:2316–2322.

Zhang, Q., Y. Li, Y. Zhang, V.E. Torres, P.C. Harris, K. Ling, and J. Hu. 2016. GTP-binding of ARL-3 is activated by ARL-13 as a GEF and stabilized by UNC-119. Sci Rep. 6:24534.

Zhang, Q., D. Nishimura, S. Seo, T. Vogel, D.A. Morgan, C. Searby, K. Bugge, E.M. Stone, K. Rahmouni, and V.C. Sheffield. 2011b. Bardet-Biedl syndrome 3 (Bbs3) knockout mouse model reveals common BBS-associated phenotypes and Bbs3 unique phenotypes. Proc Natl Acad Sci U S A. 108:20678–20683.

