## Supplementary figure legends for "ARL3 Mediates BBSome Ciliary Turnover by Promoting Its Outward Diffusion through the Transition Zone"

**Figure S1. ARL3 is highly conserved across ciliated species.** **(A)**. Sequence alignment of deduced amino acid sequences from eleven invertebrate and vertebrate ARL3 orthologues. Alignments were generated using CLC main workbench (version 6.8); the most conserved residues are shown in black, the least conserved are in red. Dashes indicate gaps introduced to optimize the alignment. **(B)**. The phylogenetic tree of ARL3 proteins from invertebrate and vertebrate species as indicated. The neighbor-joining tree was calculated using the MEGA 7 software. Branch length represents evolutionary relatedness. Accession numbers are as follows: *Bos taurus*, NP_001033656.1; *Ovis aries*, NP_001156025.1; *Homo sapiens*, AAA21654.1; *Xenopus tropicalis*, NP_AAH87495.1; *Rattus norvegicus*, NP_073191.1; *Danio rerio*, NP_001038373.1; *Mus musculus*, NP_001342162.1; *Caenorhabditis elegans*, CAB07583.1; *Trypanosoma brucei*, AAC32774.1; *Leishmania major strain Friedlin*, XP_001687211.1; and *Chlamydomonas reinhardtii*, XP_001696645.1.

**Figure S2. Characterization of the CLiP strain (LMJ.RY0420.182282) named *arl3*.** **(A)**. Agarose gel electrophoresis of the PCR products amplified from genomes of CC-5325 and *arl3* cells. The primer pair gARL3-FOR and gARL3-REV were used to amplify the *ARL3* genomic DNA of 624-bp from CC-5325 cells. A single DNA fragment of ~2.8-kb (2,848-bp) was amplified from the *arl3* cells by using the same primer pair. **(B)**. Schematic representation showing that a paromomycin-resistant gene (*AphVIII*) inserted to the fourth exon in *ARL3* gene of *arl3* cells. The boxes with colors and the lines represent the exons and introns of *ARL3* gene, respectively. **(C)**. Sequence alignment of the *ARL3* genomic DNAs (gDNA) between CC-5325 and *arl3* cells. The *arl3* cells contain a 2,284-bp *AphVIII* insertion between 520-bp and 521-bp in the *ARL3* gDNA.

**Figure S3. *Chlamydomonas* ARL3 is dispensable for ciliary assembly and IFT.** **(A)**. *arl3* is a ARL3-null mutant. Immunoblots with the affinity-purified ARL3 antiserum identified ARL3 protein with a size of approximately 20 kDa in WCE of CC-5325 but not *arl3* cells. MW stands for molecular weight. **(B)**. *arl3* cells assemble cilia with normal length. Representative phase contrast images of CC-5325 and *arl3* cells were shown (left). Scale bars: 10 µm. The *arl3* cells had full-length cilia (10.47 ± 0.82 µm, n = 20) compared to CC-5325 cells (10.57 ± 0.76 µm, n = 20). Mean lengths are listed; error bar indicates S.D. and “n” indicates the number of cilia counted. n.s.: non-significance (right). **(C)**. Immunoblots of WCE of three group cells including CC-125, CC-125::IFT43::HA::YFP, and *arl3*::IFT43::HA::YFP (left); CC-125, CC-125::IFT22::HA::YFP, and *arl3*::IFT22::HA::YFP (middle); and CC-125, CC-125::IFT38::YFP, and *arl3*::IFT38::YFP (right) probed with α-IFT43, α-IFT22, and α-IFT38, respectively. For each group of the cells, the YFP- and HA::YFP-tagged proteins of both CC-125 and *arl3* background were determined to express at the wild-type protein levels. **(D)**. Immunoblots of WCE of CC-125, *arl3*, and *arl3*::ARL3::HA::YFP probed with α-ARL3. For panels **A**, **C**, and **D**, α-tubulin we used as a loading control.

**Figure S4. Disruption of ARL3 function causes the BBSome to accumulate at the proximal ciliary region above the basal bodies.** **(A)**. *arl3*::ARL3::HA::YFP, *arl3*::ARL3^Q70L^::HA::YFP, *arl3*::ARL3^T30N^::HA::YFP, *arl3*::ARL3ΔN15^Q70L^::HA::YFP, and *arl3*::ARL3^G2AQ70L^::HA::YFP cells were stained with α-IFT46 (red) and α-HA (green). **(B)**. CC-125, *bbs8*::BBS8::YFP, and *arl3-bbs8*::BBS8::YFP cells were stained with α-IFT46 (red) and α-YFP (green). **(C)**. CC-5325, *arl3*, *arl3*::ARL3::HA::YFP, *arl3*::ARL3^Q70L^::HA::YFP, and *arl3*::ARL3^T30N^::HA::YFP cells were stained with α-BBS8 (red) and α-IFT81 (green). **(D)**. *arl3-bbs8*::ARL3::HA::YFP, *arl3-bbs8*::ARL3^Q70L^::HA::YFP, and *arl3-bbs8*::ARL3^T30N^::HA::YFP cells were stained with α-CEP290 (red) and α-YFP (green). For all four panels, phase contrast (PC) images of cells were shown. Inset shows the proximal ciliary region and the basal bodies. Scale bars: 10 µm.

**Figure S5. Knockdown of *Chlamydomonas* ARL3 does not affect ciliary assembly and IFT.** **(A)**. Representative phase contrast images of CC-125, ARL3^miRNA^, ARL3^Res-WT^, ARL3^Res-Q70L^, and ARL3^Res-T30N^ cells were shown (left). Scale bars: 10 µm. ARL3^miRNA^ (10.16 ± 1.11 µm, n = 20), ARL3^Res-WT^ (10.25 ± 0.65 µm, n = 20), ARL3^Res-Q70L^ (10.71 ± 1.07 µm, n = 20), and ARL3^Res-T30N^ (10.33 ± 0.89 µm, n = 20) cells had full-length cilia as compared to CC-125 cells (10.31 ± 0.75 µm, n = 20) (right). Mean lengths are listed; error bar indicates S.D. and “n” indicates the number of cilia counted. n.s.: non-significance. **(B)**. Immunoblots of WCE (left) and CE (right) of CC-125, ARL3^miRNA^, ARL3^Res-WT^, ARL3^Res-Q70L^, and ARL3^Res-T30N^ cells probed for the IFT-A subunits IFT43 and IFT139, the IFT-B1 subunits IFT22 and IFT70, and the IFT-B2 subunits IFT38 and IFT57. Alpha-tubulin and acetylated-α-tubulin (Ac-tubulin) were used as a loading control for WCE and CE, respectively.
