## Supplementary movie legends for "ARL3 Mediates BBSome Ciliary Turnover by Promoting Its Outward Diffusion through the Transition Zone"

**Movie S1.** TIRF imaging of ARL3::HA::YFP movement in *arl3*::ARL3::HA::YFP cilia. A frame from this movie and kymograph are shown in Fig. 1F. Play speed is real-time (15 fps).

**Movie S2.** TIRF imaging of ARL3^T30N^::HA::YFP movement in *arl3*::ARL3^T30N^::HA::YFP cilia. A frame from this movie and kymograph are shown in Fig. 2F. Play speed is real-time (15 fps).

**Movie S3.** TIRF imaging of ARL3^Q70L^::HA::YFP movement in *arl3*::ARL3^Q70L^::HA::YFP cilia. A frame from this movie and kymograph are shown in Fig. 2F. Play speed is real-time (15 fps).

**Movie S4.** TIRF imaging of ARL3ΔN15^Q70L^::HA::YFP movement in *arl3*::ARL3ΔN15^Q70L^::HA::YFP cilia. A frame from this movie and kymograph are shown in Fig. 2F. Play speed is real-time (15 fps).

**Movie S5.** TIRF imaging of ARL3^G2AQ70L^::HA::YFP movement in *arl3*::ARL3^G2AQ70L^::HA::YFP cilia. A frame from this movie and kymograph are shown in Fig. 2F. Play speed is real-time (15 fps).

**Movie S6.** TIRF imaging of BBS8::YFP movement in *bbs8*::BBS8::YFP cilia. A frame from this movie and kymograph are shown in Fig. 3E. Play speed is real-time (15 fps).

**Movie S7.** TIRF imaging of BBS8::YFP movement in *arl3-bbs8*::BBS8::YFP cilia. A frame from this movie and kymograph are shown in Fig. 3E. Play speed is real-time (15 fps).

**Movie S8.** TIRF imaging of IFT46::YFP movement in *ift46*::IFT46::YFP cilia. A frame from this movie and kymograph are shown in Fig. 5E. Play speed is real-time (15 fps).

**Movie S9.** TIRF imaging of BBS8::YFP movement in *bbs8*::BBS8::YFP cilia. A frame from this movie and kymograph are shown in Fig. 5E. Play speed is real-time (15 fps).

**Movie S10.** TIRF imaging of BBS8::YFP movement in *arl3-bbs8*::BBS8::YFP cilia. A frame from this movie and kymograph are shown in Fig. 5E. Play speed is real-time (15 fps).
