## Supplementary figures for "ARL3 Mediates BBSome Ciliary Turnover by Promoting Its Outward Diffusion through the Transition Zone"

### Slide 1
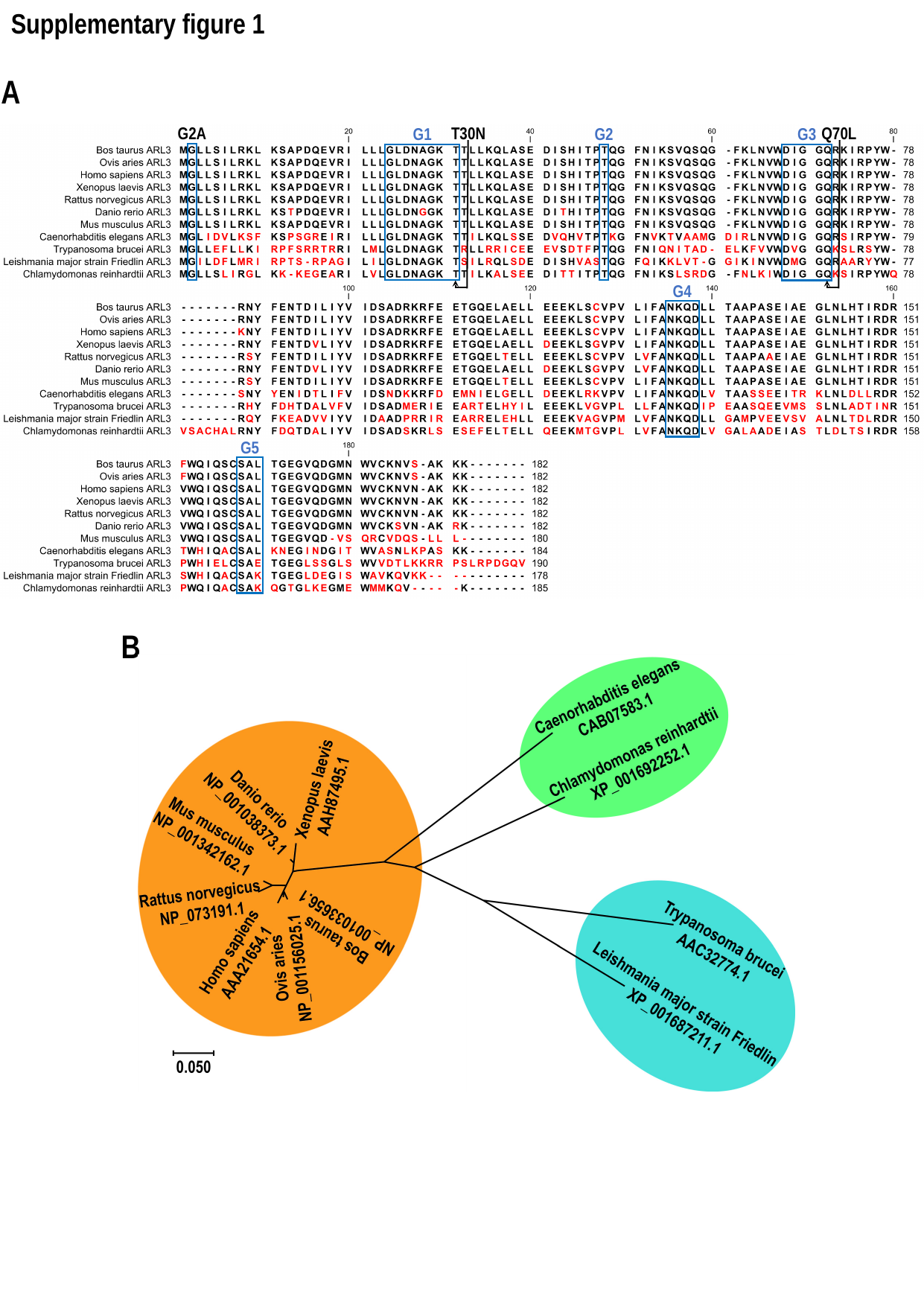

Supplementary figure 1
A
T30N
Q70L
G1
G2
G3
G4
G5
G2A
B

### Slide 2
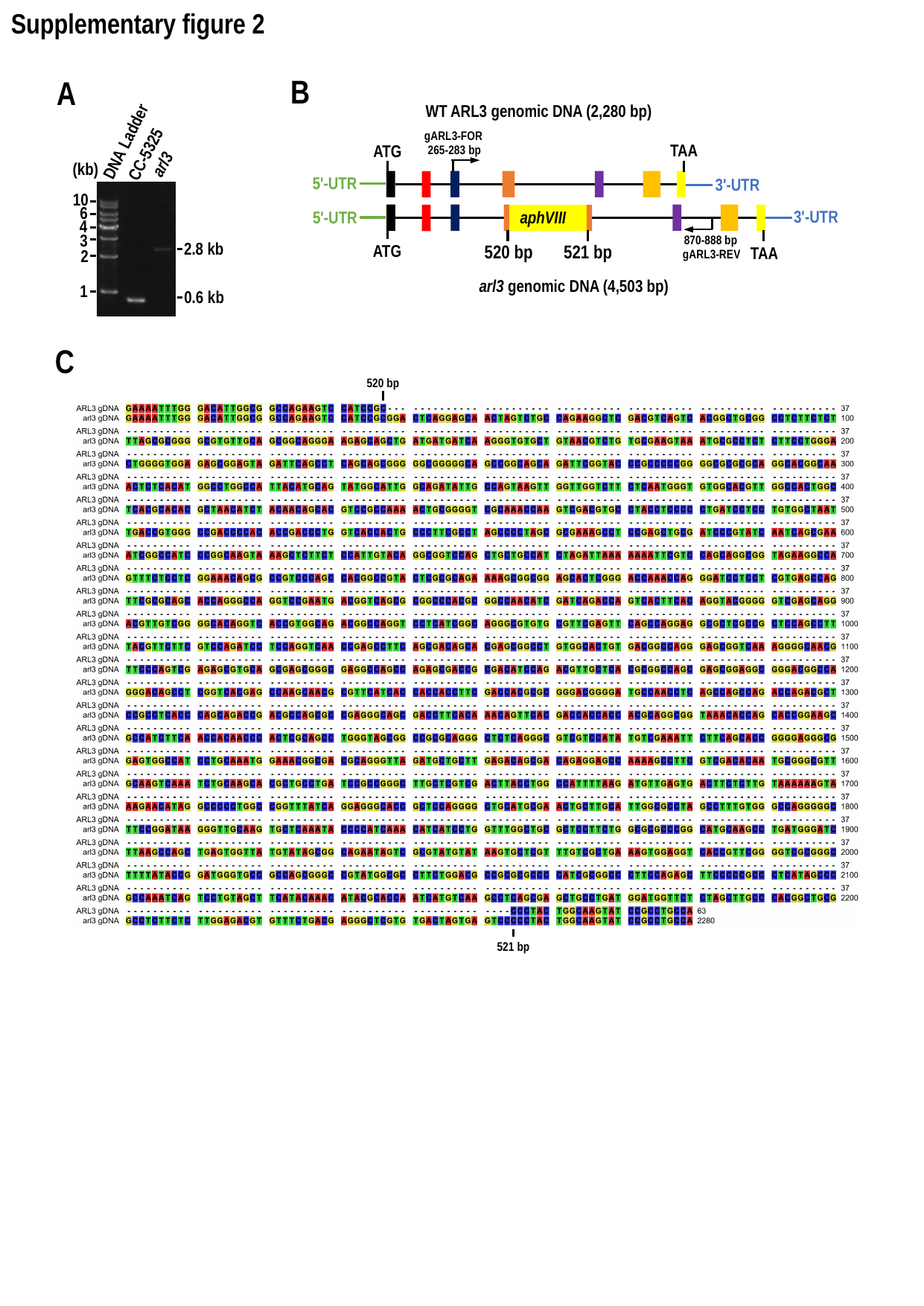

Supplementary figure 2
B
WT ARL3 genomic DNA (2,280 bp)
TAA
ATG
5'-UTR
3'-UTR
aphVIII
3'-UTR
5'-UTR
TAA
520 bp
ATG
arl3 genomic DNA (4,503 bp)
gARL3-FOR
265-283 bp
870-888 bp
gARL3-REV
521 bp
A
DNA Ladder
CC-5325
arl3
(kb)
10
6
4
3
2.8 kb
2
1
0.6 kb
C
520 bp
521 bp

### Slide 3
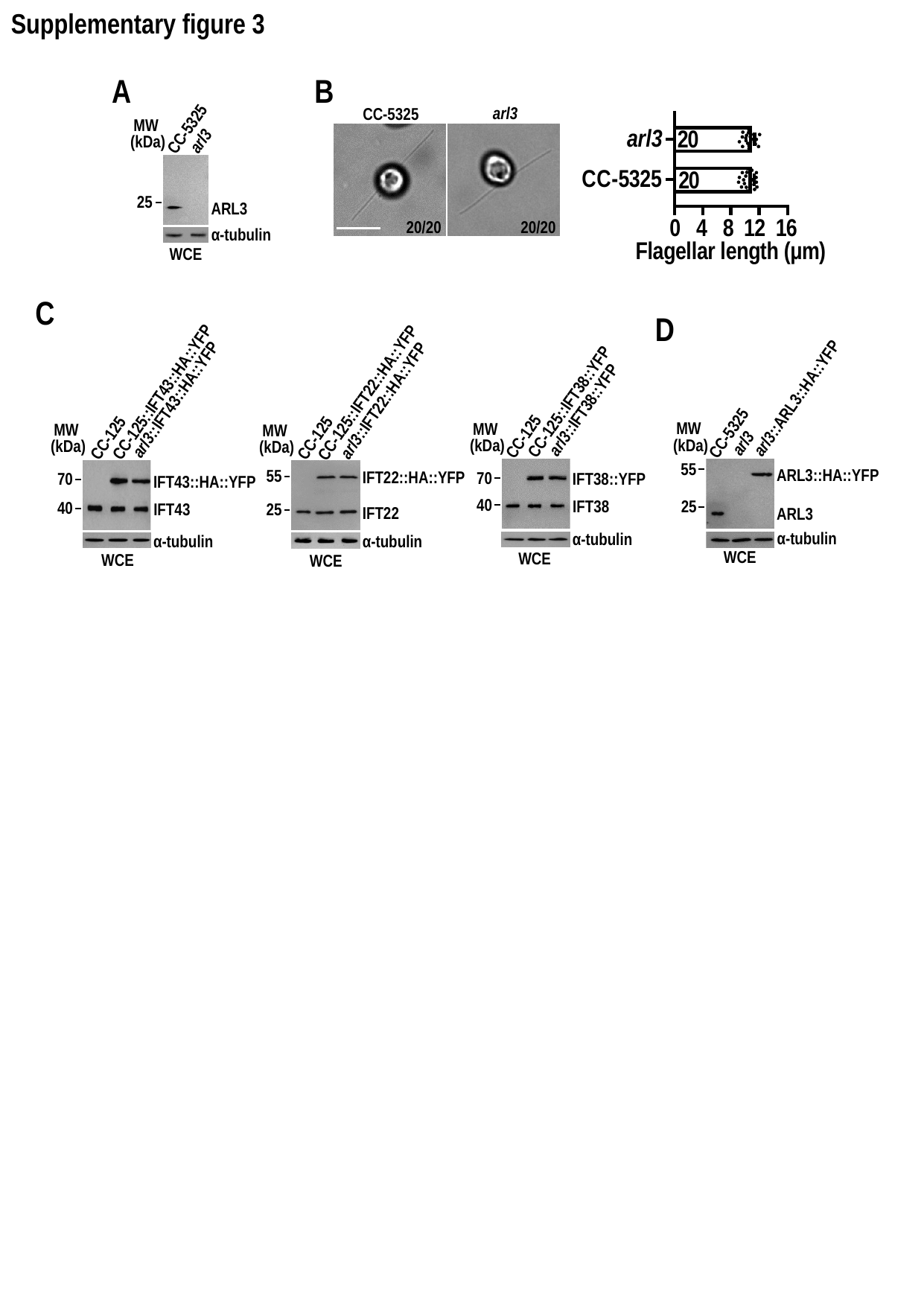

Supplementary figure 3
B
arl3
CC-5325
10 µm
20/20
20/20
A
CC-5325
MW
 (kDa)
arl3
25
ARL3
α-tubulin
WCE
C
CC-125::IFT43::HA::YFP
arl3::IFT43::HA::YFP
CC-125
MW
 (kDa)
70
IFT43::HA::YFP
40
IFT43
α-tubulin
WCE
CC-125::IFT22::HA::YFP
arl3::IFT22::HA::YFP
CC-125
MW
 (kDa)
55
IFT22::HA::YFP
25
IFT22
α-tubulin
WCE
CC-125::IFT38::YFP
arl3::IFT38::YFP
CC-125
MW
 (kDa)
70
IFT38::YFP
40
IFT38
α-tubulin
WCE
D
arl3::ARL3::HA::YFP
CC-5325
MW
 (kDa)
arl3
55
ARL3::HA::YFP
25
ARL3
α-tubulin
WCE

### Slide 4
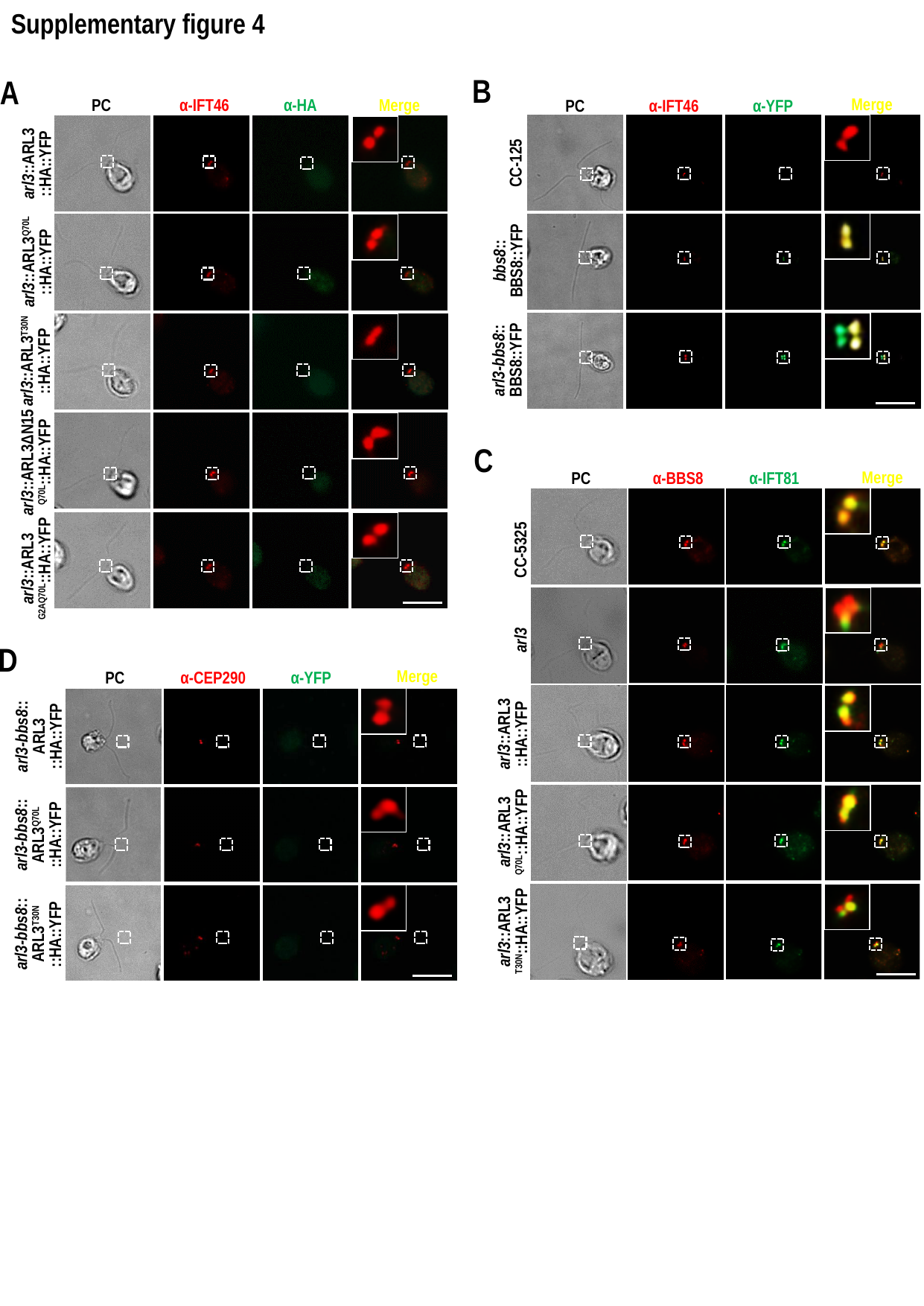

Supplementary figure 4
B
Merge
α-YFP
PC
α-IFT46
CC-125
bbs8::
BBS8::YFP
arl3-bbs8::
BBS8::YFP
A
α-HA
α-IFT46
Merge
PC
arl3::ARL3
::HA::YFP
arl3::ARL3^Q70L^
::HA::YFP
arl3::ARL3^T30N^
::HA::YFP
arl3::ARL3ΔN15
^Q70L^::HA::YFP
arl3::ARL3
^G2AQ70L^::HA::YFP
C
Merge
PC
α-BBS8
α-IFT81
CC-5325
arl3::ARL3
::HA::YFP
arl3::ARL3
^Q70L^::HA::YFP
arl3::ARL3
^T30N^::HA::YFP
arl3
D
Merge
α-CEP290
α-YFP
PC
arl3-bbs8::
ARL3
::HA::YFP
arl3-bbs8::
ARL3^Q70L^
::HA::YFP
arl3-bbs8::
ARL3^T30N^
::HA::YFP

### Slide 5
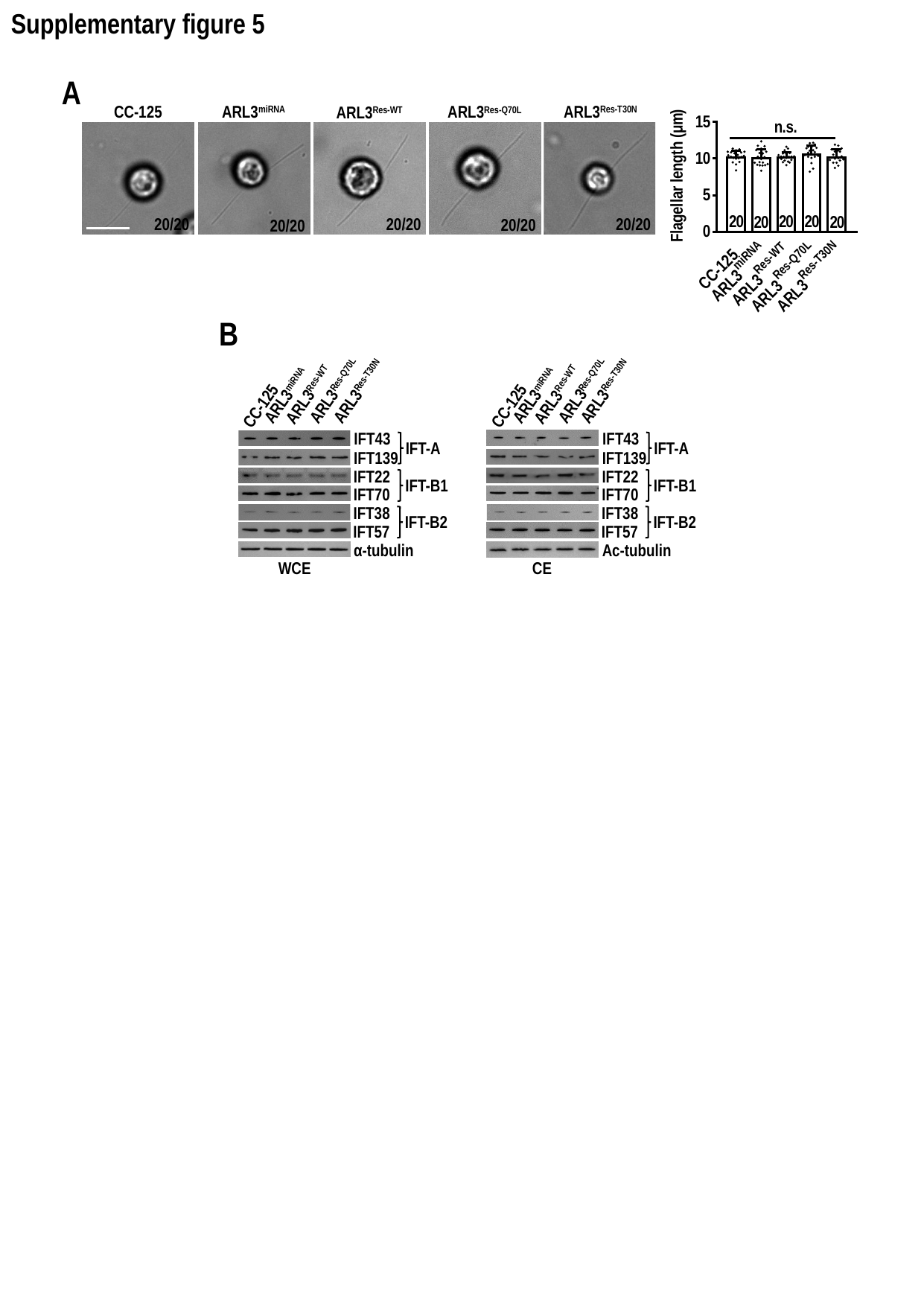

Supplementary figure 5
A
ARL3^miRNA^
CC-125
ARL3^Res-WT^
ARL3^Res-T30N^
ARL3^Res-Q70L^
20/20
20/20
20/20
20/20
20/20
B
ARL3^Res-Q70L^
ARL3^Res-Q70L^
ARL3^Res-T30N^
ARL3^Res-T30N^
ARL3^Res-WT^
ARL3^Res-WT^
ARL3^miRNA^
ARL3^miRNA^
CC-125
CC-125
IFT43
IFT43
IFT-A
IFT-A
IFT139
IFT139
IFT22
IFT22
IFT-B1
IFT-B1
IFT70
IFT70
IFT38
IFT38
IFT-B2
IFT-B2
IFT57
IFT57
α-tubulin
Ac-tubulin
WCE
CE
