## Supplementary tables for "ARL3 Mediates BBSome Ciliary Turnover by Promoting Its Outward Diffusion through the Transition Zone"

**Table S1**. Antibodies used in this study.

| **Antibody** | **Dilution**  **IB IS** | | **Origins** | **Reference or source** |
| --- | --- | --- | --- | --- |
| Anti-ARL3 | 1:250 | N/A | Rabbit | This study |
| Anti-CEP290 | 1:250 | 1:50 | Rabbit | This study |
| Anti-IFT22 | 1:1,000 | N/A | Rabbit | (Xue et al., 2020) |
| Anti-IFT38 | 1:1,000 | N/A | Rabbit | (Sun et al., 2021) |
| Anti-IFT46 | N/A | 1:100 | Rabbit | (Dong et al., 2017b) |
| Anti-IFT57 | 1:500 | N/A | Rabbit | (Dong et al., 2017b) |
| Anti-IFT43 | 1:500 | N/A | Rabbit | (Zhu et al., 2017) |
| Anti-IFT70 | 1:1,000 | N/A | Rabbit | (Dong et al., 2017b) |
| Anti-IFT81 | N/A | 1:200 | Mouse | (Fan et al., 2010) |
| Anti-IFT139 | 1:1,000 | N/A | Rabbit | (Dong et al., 2017b) |
| Anti-BBS1 | 1:1,000 | N/A | Rabbit | (Xue et al., 2020) |
| Anti-BBS4 | 1:500 | N/A | Rabbit | (Liu et al., 2021) |
| Anti-BBS5 | 1:1,000 | N/A | Rabbit | (Xue et al., 2020) |
| Anti-BBS7 | 1:500 | N/A | Rabbit | (Liu et al., 2021) |
| Anti-BBS8 | 1:500 | 1:50 | Rabbit | (Sun et al., 2021) |
| Anti-PLD | 1:1,000 | N/A | Rabbit | (Liu et al., 2021) |
| Anti-α-tubulin | 1:10,000 | N/A | Mouse | Sigma-Aldrich |
| Anti-acetylated-tubulin | 1:10,000 | N/A | Mouse | Sigma-Aldrich |
| Anti-GFP (YFP) | 1:1,000 | 1:50 | Mouse | Roche |
| Anti-HA | 1:1,000 | 1:50 | Rat | Roche |
| HRP-conjugated goat anti-rabbit IgG | 1:10,000 | N/A | Goat | The Jackson Lab. |
| HRP-conjugated goat anti-mouse IgG | 1:10,000 | N/A | Goat | The Jackson Lab. |
| HRP-conjugated goat anti-rat IgG | 1:10,000 | N/A | Goat | The Jackson Lab. |
| Alexa-Fluor 594-conjugated goat anti-rabbit IgG | N/A | 1:400 | Goat | Molecular Probes |
| Alexa-Flour 488-conjugated goat anti-mouse IgG | N/A | 1:400 | Goat | Molecular Probes |
| Alexa-Flour 488-conjugated goat anti-rat IgG | N/A | 1:400 | Goat | Molecular Probes |

Note: HRP: horseradish peroxidase. IB: immunoblotting. IS: immunostaining.

**Table S2.** *Chlamydomonas* strains used in this study

| **Name** | **Vector/background strain** | **Reference or source** |
| --- | --- | --- |
| CC-125 | Wild-type strain | CGC |
| CC-5325 | Wild-type strain | CGC |
| *arl3* | ARL3-null mutant/CC-5325 (LMJ.RY0420.182282) | CGC |
| CC-125::IFT43::HA::YFP | pBKS-gIFT43::HA::YFP-Ble/CC-125 | This study |
| CC-125::IFT22::HA::YFP | pBKS-gIFT22::HA::YFP-Paro/CC-125 | (Xue et al., 2020) |
| CC-125::IFT38::YFP | pBKS-gIFT38::YFP-Paro/CC-125 | This study |
| *arl3*::IFT43::HA::YFP | pBKS-gIFT43::HA::YFP-Ble/*arl3* | This study |
| *arl3*::IFT22::HA::YFP | pBKS-gIFT22::HA::YFP-Ble/*arl3* | This study |
| *arl3*::IFT38::YFP | pBKS-gIFT38::YFP-Ble/*arl3* | This study |
| *arl3*::ARL3::HA::YFP | pBKS-gARL3::HA::YFP-Ble/*arl3* | This study |
| *arl3*::ARL3^Q70L^::HA::YFP | pBKS-gARL3^Q70L^::HA::YFP-Ble/*arl3* | This study |
| *arl3*::ARL3^T30N^::HA::YFP | pBKS-gARL3^T30N^::HA::YFP-Ble/*arl3* | This study |
| *arl3*::ARL3ΔN15::HA::YFP | pBKS-gARL3ΔN15::HA::YFP-Ble/*arl3* | This study |
| *arl3*::ARL3ΔN15^Q70L^::HA::YFP | pBKS-gARL3ΔN15^Q70L^::HA::YFP-Ble/*arl3* | This study |
| *arl3*::ARL3ΔN15^T30N^::HA::YFP | pBKS-gARL3ΔN15^T30N^::HA::YFP-Ble/*arl3* | This study |
| *arl3*::ARL3^G2A^::HA::YFP | pBKS-gARL3^G2A^::HA::YFP-Ble/*arl3* | This study |
| *arl3*::ARL3^G2AQ70L^::HA::YFP | pBKS-gARL3^G2AQ70L^::HA::YFP-Ble/*arl3* | This study |
| *arl3*::ARL3^G2AT30N^::HA::YFP | pBKS-gARL3^G2AT30N^::HA::YFP-Ble/*arl3* | This study |
| *bbs8* | BBS8-null mutant/CC-1690 | (Sun and Pan, 2019) |
| *bbs8*::BBS8::YFP | pBKS-gBBS8::YFP-Ble/*bbs8* | This study |
| *arl3-bbs8* | ARL3- and BBS8-double null mutant | This study |
| *arl3-bbs8*::ARL3::HA::YFP | pBKS-gARL3::HA::YFP-Ble/*arl3-bbs8* | This study |
| *arl3-bbs8*::ARL3^Q70L^::HA::YFP | pBKS-gARL3^Q70L^::HA::YFP-Ble/*arl3-bbs8* | This study |
| *arl3-bbs8*::ARL3^T30N^::HA::YFP | pBKS-gARL3^T30N^::HA::YFP-Ble/*arl3-bbs8* | This study |
| *arl3-bbs8*::BBS8::YFP | pBKS-gBBS8::YFP-Ble/*arl3-bbs8* | This study |
| *ift46*::IFT46::YFP | pBKS-gIFT46::YFP-Ble/*ift46* | (Lv et al., 2017) |
| HR-YFP | pBKS-HSP70A-RBCS2-HA-YFP-Paro/CC-125 | (Dong et al., 2017a) |
| ARL3^miRNA^ | pMi-ARL3-Paro/CC-125 | This study |
| ARL3^Res-WT^ | pBKS-gARL3::HA::YFP-Ble/ARL3^miRNA^ | This study |
| ARL3^Res-Q70L^ | pBKS-gARL3^Q70L^::HA::YFP-Ble/ARL3^miRNA^ | This study |
| ARL3^Res-T30N^ | pBKS-gARL3^T30N^::HA::YFP-Ble/ARL3^miRNA^ | This study |

Note: CGC stands for *Chlamydomonas* Genetic Center.

**Table S3.** Primers used in this study

| **Name** | **Nucleotide sequence** |
| --- | --- |
| **Primers used to amplify the *aph*VIII gene insertion in ARL3 genomic DNA** | |
| gARL3-FOR | 5'-CCTCTAGAACATCACCACTATCACGC-3' |
| gARL3-REV | 5'-CCGAATTCCCAAAGCTCCGAAGCCAC-3' |
| **Primers used to clone target genes** | |
| gARL3-FOR1 | 5'-CCTCTAGAAGCTCCGACATGAGCGCC-3' |
| gARL3-REV1 | 5'-CCGAATTCCTTGACCTGCTTCATCATC-3' |
| gARL3ΔN15-FOR | 5'-CCGGATCCGTGACATGTGCTCGCAGC-3' |
| gARL3ΔN15-REV | 5'-CCGGATCCCTGTGGCGAGCACACCAG-3' |
| gIFT38-FOR | 5'-CCGGATCCATTTAGCCCCAAATTATG-3' |
| gIFT38-REV | 5'-CCGAATTCGAAGTCATTGTCATCCTC-3' |
| gIFT43-FOR | 5'-ACGCGGCCGCATGGTGTCCTACCTGGGC-3' |
| gIFT43-REV | 5'-GTGAATTCCAGCTTGATGGGCATGG-3' |
| gBBS8-FOR | 5'-GGACTAGTTGCAGCAGCAAGTAATGCAAC-3' |
| gBBS8-REV | 5'-GGAATTCCAGCATAGTGAAATGGGCC-3' |
| cARL3-FOR | 5'-GGGAATTCATGGGCCTCTTGTCGCTG-3' |
| cARL3-REV | 5'-CGCTCGAGTTACTTGACCTGCTTCATC-3' |
| cARL3ΔN15-FOR | 5'-CCGAATTCATGGCCCGCATCCTGGTC-3' |
| cARL3^G2A^-FOR | 5'-CCGAATTC ATGGCCCTCTTGTCGCTG-3' |
| cBBS1-FOR | 5'-CCGGATCCATGCTGCCATCAGTCAAG-3' |
| cBBS1-REV | 5'-CCAAGCTTTTACTCCACCTCCTCCGG-3' |
| cBBS2-FOR | 5'-CCGGATCCATGCTCGTGCCGGCCTTC-3' |
| cBBS2-REV | 5'-CCAAGCTTTCACACCGGCCCGTCGCC-3' |
| cBBS4-FOR | 5'-CAGCAAATGGGTCGGGATCCATGTCGTCATTAGCGCAG-3' |
| cBBS4-REV | 5'-TCGACGGAGCTCGAATTCTTATCACATGCCCAGCAGCT-3' |
| cBBS5-FOR | 5'-GACAGCAAATGGGTCGGGATCCATGGCTGACCTCTTTG-3' |
| cBBS5-REV | 5'-CGACGGAGCTCGAATTCTTATCACAGCACGCTCCACAG-3' |
| cBBS7-FOR | 5'-GGACAGCAAATGGGTCGGGATCCATGGAGCTGGAACTATTTC-3' |
| cBBS7-REV | 5'-GTCGACGGAGCTCGAATTCTTACTACCGCTGCCCCAGCAG-3' |
| cBBS8-FOR | 5'-GGACAGCAAATGGGTCGGGATCCATGCAGCAGCCACAGCAAG-3' |
| cBBS8-REV | 5'-GTCGACGGAGCTCGAATTCTTATCACAGCATAGTGAAATG-3' |
| **Primers used to do site-directed mutagenesis** | |
| ARL3^Q70L^-FOR | 5'-GGACATTGGCGGCCTGAAGTCCAT-3' |
| ARL3^Q70L^-REV | 5'-AGGCCGCCAATGTCCCAAATTTTC-3' |
| ARL3^T30N^-FOR | 5'-ATAACGCTGGTAAAAACACCATCCTG-3' |
| ARL3^T30N^-REV | 5'-TTTTTACCAGCGTTATCCAGTCCCA-3' |
| ARL3^G2A^-FOR | 5'-TGTGCTCGCCACAGGCCCTCTTGT-3' |
| ARL3^G2A^-REV | 5'-GCCTGTGGCGAGCACACCAGTACATG-3' |
| **Primers used to generate ARL3 miRNA construct** | |
| ARL3 miRNA-3'-UTR | 5'-ATATCAGGAAACCAAGGCGCGCTAGCTTCCTGGGCGCAGTGTTC  CAGCTGCAGTACTTAGTGAGCGATTATTTCCGTCTCGCTGATCGGCACCATGGGGGTGGTGGTGATCAGCGCTAACGGAAATAATCGCTCACTAATACTGCAGCCGGAACACTGCCAGGAGAATTC-3' |

**References**

Dong, B., H.-H. Hu, Z.-F. Li, R.-Q. Cheng, D.-M. Meng, J. Wang, and Z.-C. Fan. 2017a. A novel bicistronic expression system composed of the intraflagellar transport protein gene ift25 and FMDV 2A sequence directs robust nuclear gene expression in Chlamydomonas reinhardtii. *Applied Microbiol Biotech*. 101:4227-4245.

Dong, B., S. Wu, J. Wang, Y.X. Liu, Z. Peng, D.M. Meng, K. Huang, M. Wu, and Z.C. Fan. 2017b. Chlamydomonas IFT25 is dispensable for flagellar assembly but required to export the BBSome from flagella. *Biol Open*. 6:1680-1691.

Fan, Z.-C., R.H. Behal, S. Geimer, Z. Wang, S.M. Williamson, H. Zhang, D.G. Cole, and H. Qin. 2010. Chlamydomonas IFT70/CrDYF-1 Is a Core Component of IFT Particle Complex B and Is Required for Flagellar Assembly. *Mol Biol Cell*. 21:2696-2706.

Liu, Y.-X., B. Xue, W.-Y. Sun, J.L. Wingfield, J. Sun, M. Wu, K.F. Lechtreck, Z. Wu, and Z.-C. Fan. 2021. Bardet-Biedl syndrome 3 protein promotes ciliary exit of the signaling protein phospholipase D via the BBSome. *Elife*. 10:e59119.

Lv, B., L. Wan, M. Taschner, X. Cheng, E. Lorentzen, and K. Huang. 2017. Intraflagellar transport protein IFT52 recruits IFT46 to the basal body and flagella. *J Cell Sci*. 130:1662-1674.

Sun, L., and J. Pan. 2019. Bardet-Biedl syndrome protein-8 is involved in flagellar membrane protein transport in Chlamydomonas reinhardtii. *Sheng Wu Gong Cheng Xue Bao*. 35:133-141.

Sun, W.-Y., B. Xue, Y.-X. Liu, R.-K. Zhang, R.-C. Li, W. Xin, M. Wu, and Z.-C. Fan. 2021. Chlamydomonas LZTFL1 mediates phototaxis via controlling BBSome recruitment to the basal body and its reassembly at the ciliary tip. *Proceedings of the National Academy of Sciences*. 118:e2101590118.

Xue, B., Y.-X. Liu, B. Dong, J.L. Wingfield, M. Wu, J. Sun, K.F. Lechtreck, and Z.-C. Fan. 2020. Intraflagellar transport protein RABL5/IFT22 recruits the BBSome to the basal body through the GTPase ARL6/BBS3. *Proc Natl Acad Sci U S A*. 117:2496-2505.

Zhu, B., X. Zhu, L. Wang, Y. Liang, Q. Feng, and J. Pan. 2017. Functional exploration of the IFT-A complex in intraflagellar transport and ciliogenesis. *PLOS Genet*. 13:e1006627.
